# Metagenomic global survey and in-depth genomic analyses of *Ruminococcus gnavus* reveal differences across host lifestyle and health status

**DOI:** 10.1101/2024.06.27.600998

**Authors:** S. Nooij, N. Plomp, I.M.J.G. Sanders, L. Schout, A.E. van der Meulen, E.M. Terveer, J.M. Norman, N. Karcher, M.F. Larralde, R.H.A.M. Vossen, S.L. Kloet, K.N. Faber, H.J.M. Harmsen, G.F. Zeller, E.J. Kuijper, W.K. Smits, Q.R. Ducarmon

**Affiliations:** Leiden University Center of Infectious Diseases (LUCID), Leiden University Medical Center, Leiden, The Netherlands; Center for Microbiome Analyses and Therapeutics, Leiden University Medical Center, Leiden, The Netherlands; Department of Medical Microbiology and Infection Prevention, University of Groningen, University Medical Center Groningen, Groningen, The Netherlands; Department of Gastroenterology and Hepatology, Leiden University Medical Center, Leiden, The Netherlands; Vedanta Biosciences, Inc., Cambridge, Massachusetts, USA; Structural and Computational Biology Unit, European Molecular Biology Laboratory, 69117 Heidelberg, Germany; Leiden Genome Technology Center, Leiden University Medical Center, Leiden, The Netherlands; Department of Gastroenterology and Hepatology, University of Groningen, University Medical Center Groningen, Groningen, The Netherlands

**Keywords:** inflammatory bowel disease, metagenome-assembled genomes, PacBio Hi-Fi, strain variation, Crohn’s disease

## Abstract

*Ruminococcus gnavus* is a highly prevalent gut bacterium (present in >90% of healthy individuals), of which increased abundance is associated with chronic inflammatory diseases, most notably Crohn’s disease. Nevertheless, its global distribution has not been investigated and little is known about intraspecies genomic variation. Through a large-scale survey of 12,791 gut metagenomes, we recapitulated known associations with metabolic diseases and inflammatory bowel disease. We uncover a higher prevalence and abundance of *R. gnavus* in Westernized populations and observe relative abundances of up to 83% in newborns and infants. Next, we built a collection of existing and newly cultured *R. gnavus* isolates (N = 45) from both healthy individuals and Crohn’s disease patients and subjected these to PacBio circular consensus sequencing to greatly expand the number of complete *R. gnavus* genomes. Analysis of these genomes as well as publicly available high quality draft genomes (total > 300 genomes) revealed multiple clades which separated Crohn’s-derived isolates from healthy-derived isolates. Functional analyses of genes predicted to constitute *R. gnavus* virulence factors could not explain this separation. Bacterial GWAS revealed that Crohn’s-derived isolates were enriched in genes related to mobile elements and putative mucin foraging. Together, we present one of the largest complete genome collections of any commensal gut microbe and provide novel biological insights into the global distribution and genomic variation of *R. gnavus*.

## INTRODUCTION

The human gut microbiome is a topic of intense research interest and many bacterial species have been associated with specific diseases^1^. One such species is *Ruminococcus gnavus*, for which associations with human health have been reported in the context of various ailments^2–7^. Officially, its taxonomic status has been revised and *R. gnavus* is now member of the genus *Mediterraneibacter*^8^. *R. gnavus* is a non-spore forming Gram-positive member of the bacterial phylum Bacillota (formerly Firmicutes) and was first described in 1976^9^. It is considered a prevalent member of the gut microbiome (present in > 90% of healthy European and North-American adults) and its median relative abundance is reported to be approximately 0.1% – 0.3%, although it should be noted that these estimates were based on small and geographically restricted studies^10,11^.

In microbiome association studies, increases in *R. gnavus* relative abundance have consistently been linked to diseases including metabolic syndrome, type 2 diabetes mellitus and Crohn’s disease (CD, a form of inflammatory bowel disease (IBD))^2,3,12^. Furthermore, its relative abundance increased concomitantly with symptomatic flares in CD, where it reached up to 69.5% of the gut microbiome^2^. While it remains unknown if *R. gnavus* causally contributes to disease development or whether the increased abundance is a result of the changing intestinal environment, several molecular mediators have been identified that potentially contribute to disease. For instance, the cell-surface exposed polysaccharide glucorhamnan has been described as pro-inflammatory, with a strain-dependent effect, depending on whether the *R. gnavus* isolate carried a capsular polysaccharide that promoted a more tolerogenic response^13,14^. However, these observations are limited by the fact that they were made using one or few isolates and strain variation remains underexplored in many gut microbes, including *R. gnavus*.

Not only mechanistic, but also genomic studies of *R. gnavus* to date have suffered from these limitations. One study divided *R. gnavus* into two clades based on genome sequences and noted that one was enriched in IBD patients^2^. However, this study was limited by a low number of draft isolate genomes (N = 11) and a scarcity of knowledge on experimentally verified virulence factors of *R. gnavus* at the time^13–18^. A more recent study based on 152 draft genomes identified three major lineages, but genomes of different host organisms were mixed and this study did not investigate associations of genetic features with metadata^19^. Therefore, an important outstanding question remains whether proposed *R. gnavus* virulence factors are enriched in IBD-derived isolates, or whether different genes and functions could separate IBD-derived *R. gnavus* isolates from controls.

Here, we surveyed global *R. gnavus* prevalence and abundance across thousands of gut metagenomes to provide a more nuanced picture across human lifespan, different lifestyles, and disease, thereby revealing striking differences. Next, through extensive culturing efforts we established a collection of 45 *R. gnavus* isolates and applied PacBio circular consensus sequencing (CCS) to generate complete genomes. Complementing this unique collection with publicly available (short-read draft) genomes allowed us to perform large-scale comparative genomics at both the level of phylogeny and predicted gene functions.

## RESULTS

### Intestinal colonization with *R. gnavus* is associated with age, health, geography and lifestyle

In order to provide a nuanced view of *R. gnavus* prevalence and abundance across health and disease, geography, and lifestyle, we screened 12,791 publicly available metagenomes from all over the world with manually curated metadata (Figure 1)^20^. We observed *R. gnavus* in 50.58% of all included subjects and the prevalence in 9,126 healthy individuals was 43.09% (Figure 1A). As *R. gnavus* has been robustly associated with disease, especially with metabolic disease and IBD^2,3^, we compared *R. gnavus* prevalence and abundance between patients with these diseases and healthy subjects (or asymptomatic control subjects) in a meta-analysis. *R. gnavus* was approximately 1.6 times more prevalent in IBD patients (70.2%; Chi-square, adjusted p = 4.6 × 10^−40^, odds ratio (OR [95% confidence interval]) = 3.1 [2.6-3.7]), 1.3 times more with hypertension (58.0%; p.adj = 0.0024, OR = 1.8 [1.3-2.5]), 1.5 times with type-2 diabetes (T2D; 62.9%; p.adj = 1.9 × 10^−11^, OR = 2.2 [1.8-2.8]), and 2.2 times with atherosclerotic cardiovascular diseases (ACVD; 96.2%; p.adj = 3.9 × 10^−52^, OR = 33.4 [17.6-74.1]) compared to healthy subjects. Furthermore, the relative abundance of *R. gnavus* was also higher in these conditions as compared to healthy (Figure 1B; healthy: median = 0%, 1^st^−3^rd^ quartile [0-0.08%]; IBD: median = 0.11% [0-1.04%], t-test, p.adj = 2.3 × 10^−48^; hypertension: median = 0.01% [0-0.07%], p.adj = 0.007, T2D: p.adj = 2.3 × 10^−11^; ACVD: median = 0.78% [0.09-3.14%], p.adj = 7.5 × 10^−78^). Together, we thus recapitulated that *R. gnavus* occurs more frequently and in higher abundances in the gut microbiome of patients suffering from IBD, hypertension and T2D. Additionally, our analysis uncovered a striking novel enrichment in ACVD, which had the highest prevalence and abundance of any disease group.

**Figure 1.**
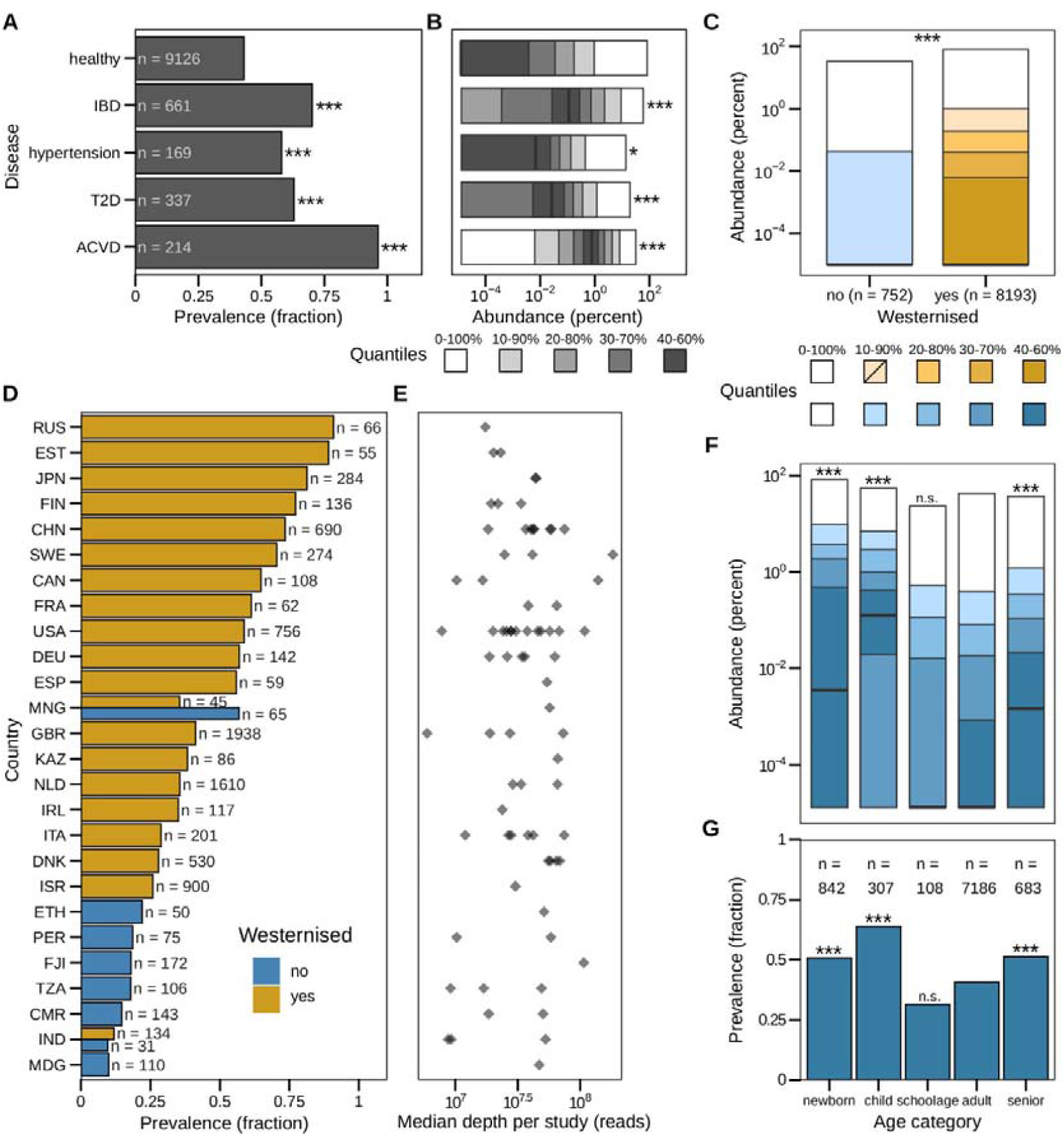
Intestinal colonization with *R. gnavus* is associated with age, health, geography, and lifestyle. (A) We queried the public resource curatedMetagenomicData for relative abundances of *R. gnavus* in human stools to conduct a meta-analysis of global prevalence and abundance. Prevalence is shown as fraction of subjects with *R. gnavus* abundance > 0, grouped by selected health conditions. IBD: inflammatory bowel diseases, T2D: type-2 diabetes, ACVD: atherosclerotic cardiovascular diseases. Each disease is compared to healthy (*** p < 0.001). (B) Relative abundance of *R. gnavus* in the same groups as (A), shown as quantile plots, using quantiles ranging from 0 to 100% in increments of 10 with the median shown as a thick black line and quantiles closer to the median shown as darker shades of the same color (see Methods). Each disease is compared to healthy (* p < 0.05, *** p < 0.001). (C) Comparison of *R. gnavus* abundance between healthy people from Westernized and non-Westernized societies as quantile plot. The p-value was calculated using a t-test (*** p < 0.001). (D) Prevalence of *R. gnavus* grouped per country and colored by Westernization, only showing results from countries from which at least 50 samples were collected. (Countries are abbreviated by ISO 3166-1 alpha-3 codes.) (E) Sequencing depth control per country (same as (D)). Each diamond represents a study that collected samples from the corresponding country. Sequencing depth is shown as median number of reads generated per country in the study. (F) Relative abundance of *R. gnavus* in different age categories (newborn: < 1 year, child: 1-11 years, school age: 12-18 years, adult: 19-65, senior: 65+ years) shown as quantile plots. Age categories are listed in (G). Each age category is compared to adult (*** p < 0.001, n.s. not significant). (G) Prevalence of *R. gnavus* among different age categories. Each category is compared to adult (*** p < 0.001, n.s. not significant). In (B), (C), and (F), a pseudocount of 1.3 × 10^−5^ is added to all abundances to enable visualization on a logarithmic scale. See also Figure S1.

Subsequently, we investigated prevalence and relative abundance of *R. gnavus* across countries (Figures 1C, 1D and S1A). In this analysis, we included only healthy individuals to account for the possibility of confounding by diseases such as IBD and metabolic disease. We observed large differences in prevalence, which ranged between 10-90% across countries (overall median: 41%) and mean relative abundance per country ranged between 0.0078-4.05% (overall mean = 0.67% ±3.20 standard deviation; Figure S1A). This variation could be partly explained by Westernization status; this binary classification of Westernized / non-Westernized lifestyles is based on, among others, access to medical care and pharmaceuticals, livestock exposure and diet^21^. Westernized individuals had higher prevalence and abundance of *R. gnavus* compared to non-Westernized individuals (Figure 1C-E; prevalence: Chi-square p = 2.6 × 10^−47^). As these data were generated in multiple studies, we cannot exclude effects of technical differences (e.g., DNA extraction method). To partially check for this, we investigated sequencing depth and found that higher prevalence and abundance were not the result of higher sequencing depth in Westernized countries as non-Western samples were sequenced deeper (Figure S1B; t-test p = 6.9 × 10^−20^). These differences hold true for any 10% quantile of sequencing depth (Figure S1C, Methods). We also checked for possible correlations between sequencing depth and *R. gnavus* abundance and found a weakly negative correlation in both Westernized and non-Westernized metagenomes (Figure S1D). In conclusion, *R. gnavus* colonization is vastly different between countries, and Westernization (lifestyle) may be a major factor contributing to these differences.

We noted extremely high *R. gnavus* abundance values in healthy people, up to a relative abundance of 83%. Metagenomes with the highest abundances were often samples collected from newborns and children up to age 2, most of whom were recorded not to have received antibiotics. This motivated a further analysis of age-related patterns of *R. gnavus* colonization (Figure 1F)^20^. *R. gnavus* abundances were higher in newborns (t-test, p.adj = 3.7 × 10^−45^) and children up to 11 years old (p.adj = 4.1 × 10^−34^) as compared to adults, and abundances were also higher in seniors (65-92 years old) than in adults (p.adj = 3.7 × 10^−11^). We observed similar patterns regarding *R. gnavus* prevalence (Figure 1G), where newborns (Chi-Square, p.adj = 2.5 × 10^−7^), children aged 1-11 (p.adj = 1.3 × 10^−14^) and seniors (p.adj = 8.1 × 10^−7^) were more likely to carry *R. gnavus* than adults. While the high prevalence and abundance in newborns and infants has previously been reported^22^, we here present the largest and most comprehensive analysis on this topic, which supports the notion that *R. gnavus* is a common and predominant inhabitant of the infant gut.

In summary, colonization with *R. gnavus* appears to be dynamic across the lifespan in healthy individuals, with the highest abundances observed in newborns. While these metagenomic analyses provide important insight into the global distribution of *R. gnavus*, in-depth genomic analyses are required to investigate whether genomic content differs across described factors such as disease and geography.

### Newly generated complete genomes have superior assembly characteristics and cover phylogenetic diversity

For our large-scale genomic analysis of *R. gnavus*, we first established an isolate collection through extensive culturing efforts and by collecting available isolates, from which we sequenced the genome of 45 isolates using PacBio circular consensus sequencing (CCS) to yield complete, circular genomes and potential extrachromosomal elements (Figure 2, Methods). We next complemented these with 208 available MAGs for which sufficient metadata could be retrieved and short-read genome data of an additional 79 isolates (Methods). Of note, given the relative infancy of assembly algorithms for PacBio CCS data of microbial genomes, we performed a mini-benchmark of five long-read *de novo* assemblers. This indicated clear differences in number of contigs generated, as well as in runtime and memory usage between assemblers (Figure S2, Methods). Comparing the quality of these genomes, we observed that MAGs were worse in every aspect of genome assembly when compared to isolate assemblies (Figure 2A). While total length and number of genes were lower for MAGs as expected, GC content clearly differed between MAGs and isolate genomes, suggesting that current MAG binning techniques may fail to capture AT-rich regions. We further observed that isolates that underwent PacBio CCS were often assembled into single circular contigs, in contrast to a mean of 107 (±58.4 standard deviation) contigs per short-read isolate genome. Additionally, we found four circular extrachromosomal elements predicted to be plasmids with 99.9% confidence (Figure 2A, 2B, Figure S3), demonstrating the added value of PacBio CCS. These four putative plasmids comprise two different large sequences of 191kb and 164kb, which derived from two distinct isolates from healthy individuals (i.e., QRD006, QRD009 and QRD010 contain one plasmid, QRD011 the other), and have not been described in *R. gnavus* to date. The plasmids are modular and highly related, that is, they are identical except for one gene cluster that is missing from the shorter 164kb plasmid (Figure S3A).

**Figure 2.**
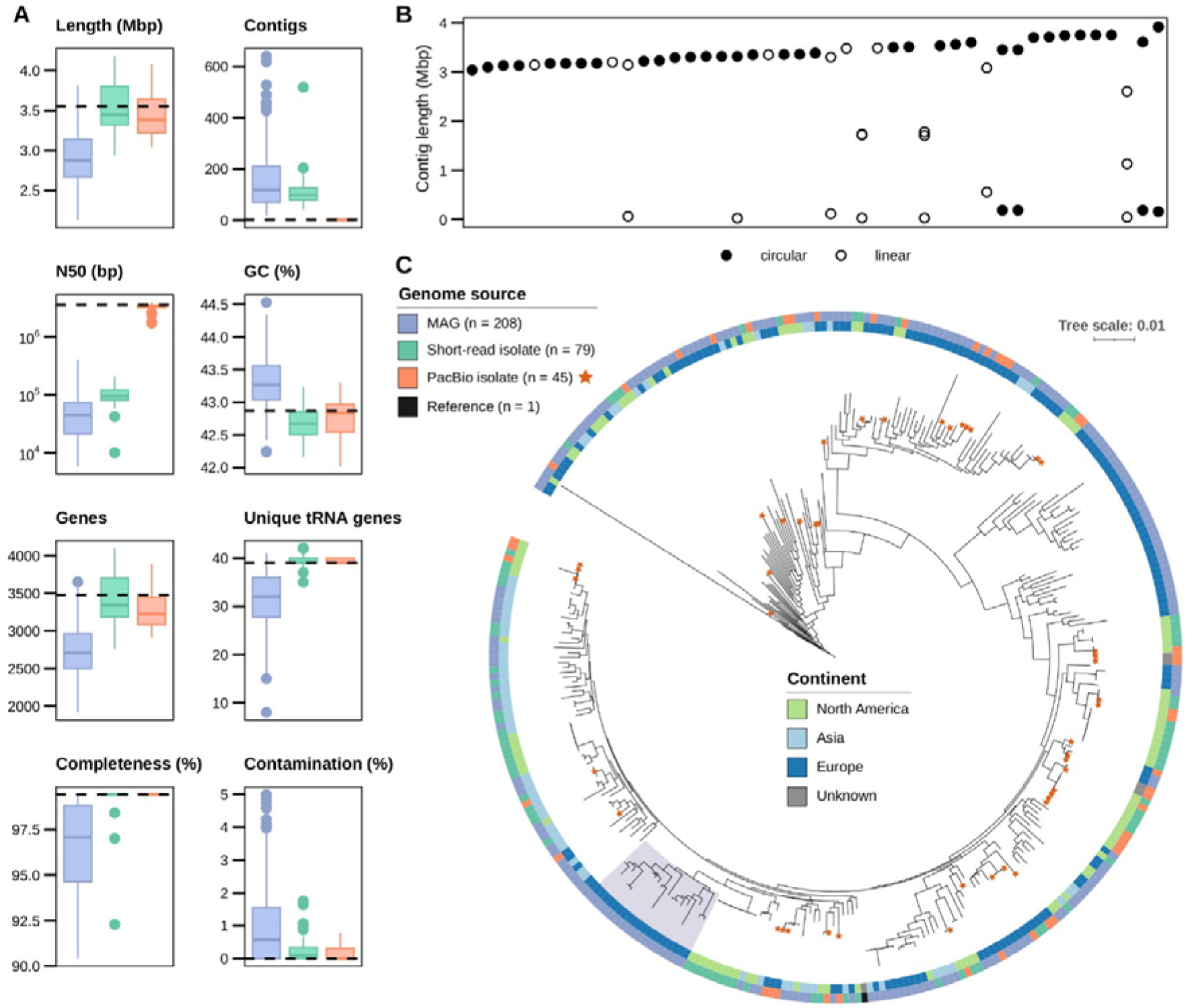
Newly generated complete genomes have superior assembly characteristics and cover phylogenetic diversity. (A) We collected both publicly available short-read-based genomes from isolates and metagenome-assembled genomes (MAG), as well as long-read genomes generated from isolates in this study using PacBio HiFi sequencing and compared them to the one reference genome from NCBI GenBank (accession number GCF_009831375.1). Assembly statistics of each group of genomes are compared to the reference genome, shown as dashed line. Thick lines indicate medians, boxes represent first and third quantile and whiskers indicate the rest of the data excluding outliers; outliers are shown as separate dots. Color legend is shared with (C). (B) Length and circularity of *de novo* assembled contigs from PacBio HiFi reads. (C) Maximum likelihood phylogenetic tree based on concatenated core genes. Each genome is annotated with its corresponding genome source and continent of origin. Stars mark genomes sequenced with PacBio newly added in this work. The gray shaded area marks the infant-associated clade that contains 8/10 MAGs with flagellum genes. See also Figure S2, S3, S4, S5 and S6.

They do not contain evident predicted antibiotic resistance or virulence genes (Methods). The plasmids are likely conjugative or mobilizable based on identified putative transposase genes which is consistent with their geographically distinct origins (USA and Japan). The plasmids contain a putative ParABS segregation system, annotated as ‘Soj’ (ParA) and ‘ParB domain containing protein’ (ParB). A key feature is a (hypothetical) non-ribosomal protein synthesis (NRPS) cluster with no known homologs (Figure S3B). However, upstream of it we identified with moderate confidence a transcription factor binding site for CatR, an H_2_O_2_-responsive repressor.

Leveraging our large genome collection, we then investigated the phylogenetic diversity of *R. gnavus* (Figure 2C). This revealed no continent or genome source-specific clustering, but importantly, demonstrated that our *R. gnavus* isolate collection captures the full breadth of phylogenetic diversity across the tree (Figure 2C).

Using PacBio’s capability to identify methylated DNA, we identified 18 motifs associated with m^4^C and 32 motifs associated with m^6^A (Figure S4), greatly expanding our knowledge on methylation in this organism. One methylated motif (VNNVNCTGVNCAN) appeared conserved across all isolates but one, while the rest of the motifs was isolate-specific and could not be linked to phylogeny (Figure S4, Table S4).

### *R. gnavus* motility possibly restricted to infant-derived strains

In order to characterize the functional capacity of *R. gnavus*, we annotated our genomes with functional orthologs, modules and pathways (from KEGG^23^) and used linear modelling to identify associations between microbial functions and metadata. Using this methodology, we observed flagellum biosynthesis exclusively in newborns and infants up to 1 year of age, and this association was also statistically significant (p = 0.008). We further investigated flagellum biosynthesis together with chemotaxis, as these are functionally closely related, and found both pathways in ten out of 333 genomes. These ten genomes are all MAGs originating from newborns and infants up to 1 year of age (Figure S5A) and contained (almost) complete operons (Figure S5B). To ensure this finding was not a technical assembly artefact, we traced the origin of these genomes, which revealed that these MAGs derive from infants sampled in three studies and five geographically separated locations (Estonia, Finland, Italy, Russia, and Sweden). Eight out of ten genomes with flagellum genes belong to a phylogenetic clade that is associated with newborns and infants (17/19 genomes in that clade derive from infants of 1 year old or younger; Figure 2C, clade highlighted in gray), suggesting that motility might be associated with a specific infant-associated clade of *R. gnavus*. The absence of isolates in this clade precludes experimental verification of flagellum functionality, but strain differences in flagella and motility have been described^9^.

We also screened all genomes for antibiotic resistance genes and found that resistance against tetracycline is the most common among *R. gnavus* (75/125 isolates; Figure S6). *In vitro* validation confirmed that at least isolates containing the *tet(O)* and/or *tet(40)* genes, which account for the majority of the observed tetracycline resistance determinants, indeed have increased minimum inhibitory concentrations compared to isolates without *tet* gene (Table S2).

### Genomic differences between isolates from healthy and Crohn’s indicates a Crohn’s-specific subspecies

To evaluate whether CD-derived *R. gnavus* isolates genomically differ from healthy-derived isolates, we first placed our genomes into a core genome-based phylogenetic tree (Figure 3A). As this tree contains practically identical isolates derived from the same person, we also constructed a tree of deduplicated genomes to facilitate statistical testing (Figure S7). This revealed three main clades with a strong enrichment of Crohn’s-derived isolates in the two more basal clades (Fisher’s exact test, p = 0.00037, OR = 12.1 [2.5-69.8]). As our phylogenetic tree was reconstructed from only the core genome, we next performed whole-genome ANI analysis and accessory genome comparisons to also assess differences in the other genomic loci, which resulted in a highly similar clustering (Figure 3A, 3B). As all *R. gnavus* genomes included here share at least 95% similarity with one another, which is often considered the species boundary^24,25^, we consider that these clades represent subspecies. Subsequently, we clustered all core genomes using Principal Coordinate Analysis (PCoA) of the phylogenetic distances and k-means clustering (Figure S7) and found a clear separation between the Crohn’s-associated clades and the healthy-associated clade (Chi-square, p = 1.25 × 10^−6^, OR = 5.8 [2.7-12.9]). Together, these results demonstrate that at *R. gnavus* isolates from CD patients are genomically distinct from isolates from healthy controls based both on their core and accessory genome. The phylogeny indicates that most healthy-derived isolates form a monophyletic subspecies clade, while the CD isolates appear polyphyletic and may be categorized into multiple groups.

**Figure 3.**
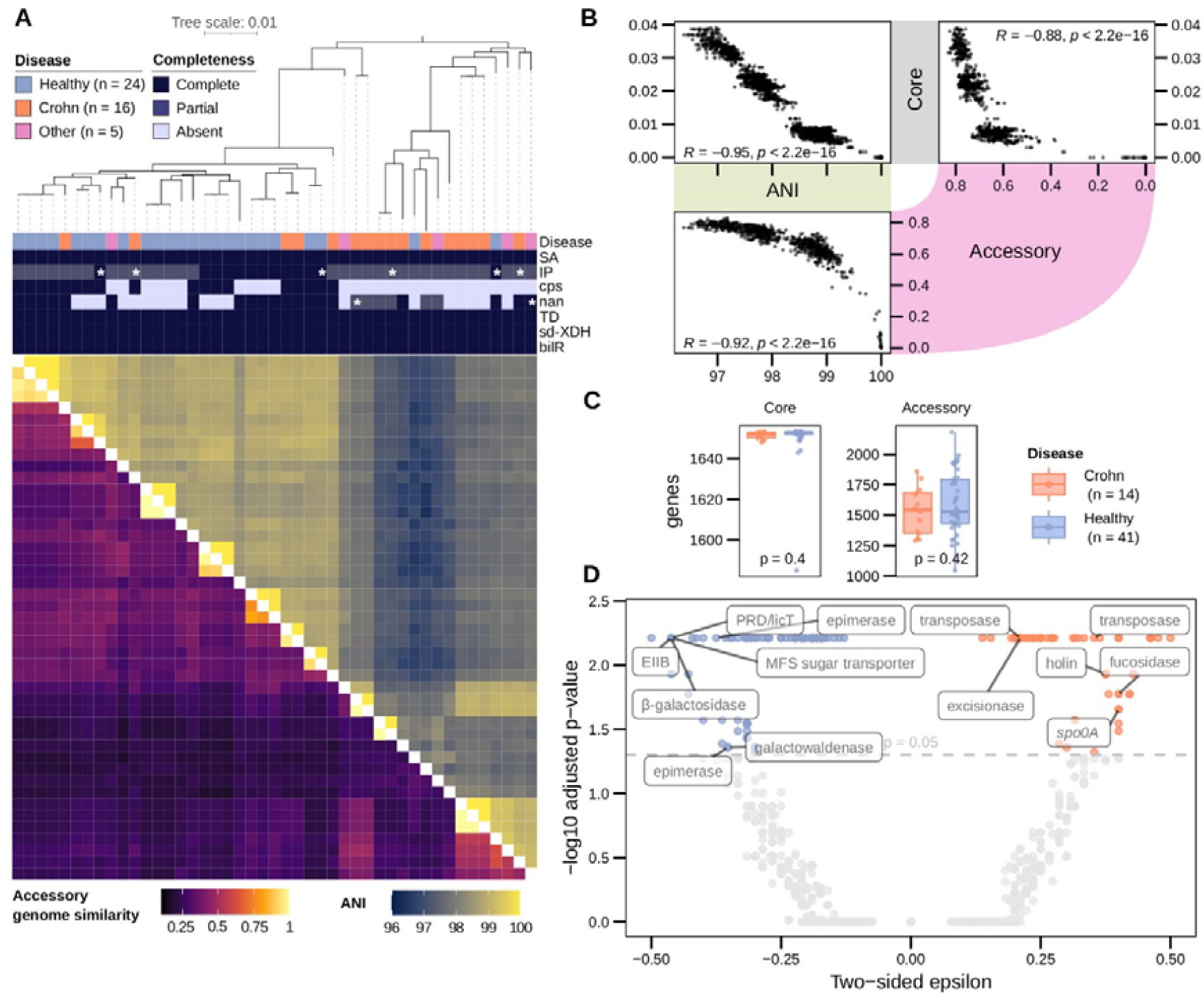
Genomic differences between isolates from healthy and Crohn’s indicates a Crohn’s-specific subspecies. (A) Using our newly generated PacBio genomes, we compared genomes of isolates from healthy people to isolates from CD patients. Maximum likelihood phylogenetic tree of PacBio isolate genomes using concatenated core genes, with annotation of disease status and genes and gene clusters described previously in literature. Asterisks indicate gene clusters from genomes that are highlighted in Figure S6. Below are heatmaps of pairwise average nucleotide identity (ANI) and accessory genome similarity (calculated as 1 / binary distance). SA: superantigen (2 genes), IP: inflammatory polysaccharide (23 genes, ‘partial’ = 20 or 21 genes), cps: capsular polysaccharide (20 genes), nan: sialic acid metabolic cluster (11 genes, ‘partial’ = 6 genes), TD: tryptophane decarboxylase (1 gene), sd-XHD: selenium-dependent xanthine dehydrogenase (1 gene), bilR: bilirubin reductase (1 gene). (B) Comparison of genome comparison metrics core genome phylogenetic distance, average nucleotide identity and accessory genome binary distance tested with Spearman correlations. (C) Comparison of core and accessory genome size between deduplicated isolate genomes with a CD or healthy phenotype, derived from short-read or long-read sequencing. (D) We compared accessory genomes of isolates from healthy people and CD patients using a bacterial GWAS to identify genes associated with disease phenotype. Results are expressed as false discovery rate-adjusted p-value (using the Benjamini-Hochberg correction) and epsilon, which is a measure of association strength between phenotype and genotype based on the (maximum likelihood) phylogenetic tree. The gray dashed line indicates a p-value of 0.05, anything above the line is considered statistically significant. Positive values of epsilon correspond to an enrichment in CD and negative epsilon values are associated with a healthy host phenotype. See also Figure S7, S8, S9 and S10.

### Host phenotypes cannot be explained by previously identified putative virulence factors in *R. gnavus*

We next tested whether previously suggested *R. gnavus* virulence factors could explain the association with CD (Figure S7). First, we observed that four genes or gene clusters (superantigens, tryptophane decarboxylase, bilirubin reductase and selenium-dependent xanthine dehydrogenase) were present in all complete *R. gnavus* genomes and are therefore part of the core genome (Figure 3A). While we saw variation in several other gene clusters (glucorhamnan-producing gene cluster, Fisher’s exact test, p = 0.19; and the *nan* gene cluster, p = 0.35), only one, namely the capsular polysaccharide gene (*cps*) cluster was associated with the distinction and was detected exclusively in isolates from the healthy-associated clade (p = 0.0008). In conclusion, only the *cps* cluster, that leads to a more tolerogenic immune response^14^, could distinguish host phenotype groups.

### Genomic architecture of gene cluster producing the proinflammatory polysaccharide glucorhamnan reveals genomic variations

Previous studies have highlighted the relevance and genomic architecture of the gene cluster producing inflammatory glucorhamnan based on complete, intermediate, or limited short-read coverage^26^. Here, we re-examined in our diverse collection of complete genomes if these clusters derive from the same genomic locus and are likely to be homologous (Figure S8). Compared to the isolate in which the gene cluster was experimentally verified (QRD039 = RJX 1121)^13^, we saw variations in multiple genes, including several glycosyltransferases (Figure S8A). We observed 13 out of 45 long-read genomes to have the complete original cluster as identified in RJX1121, while 30 genomes had 20/23 genes as annotated in NZ_AAYG02000032.1 and two had 21/23 genes (those with 20 or 21 hits are subsequently called ‘partially complete’)^13,26^. These genomes lacked the same genes: a glycosyltransferase (RUMGNA_03519; present in the two genomes with 21 genes found), a transporter (RUMGNA_03522) and a polyphosphoglycerol synthesis gene (RUMGNA_03523). These partially complete cluster variants lack the genes in positions that were reported to have low coverage and we think they are therefore the same as those described in Sorbara *et al*., 2020 as ‘intermediate coverage’. To elucidate whether these genomes contain a truly different gene cluster at a different genomic location, the flanking genes were determined to map the genomic neighborhood. All investigated genomes had the same neighboring genes, thereby revealing a conserved genomic locus. (The 3’ and 5’-flanking genes are annotated as ‘HPr family phosphocarrier protein’ and ‘glutamine-fructose-6-phosphate transaminase’.) By closer inspection of the genomic loci, we found that the operon lacking RUMGNA_03519, RUMGNA_03522 and RUMGNA_03523 had other genes inserted instead (Figure S8A). Moreover, the variability at protein level compared to the reference gene (30-70% identity) suggests that this whole locus may be subject to positive selection or adaptation pressure. Nevertheless, based on similarity in genomic architecture we expect that all these strains still produce polysaccharides, although it remains to be established whether all of them mediate pro-inflammatory effects.

A similar comparative genomics analysis for the *nan* gene cluster, responsible for releasing 2,7-anhydro-Neu5Ac from mucin^18^, showed some genomes with *nan*-like genes in a different locus (Figure S8B). All these alternative *nan*-like clusters had the same genomic architecture, which importantly lacked the *nanH* (intramolecular trans sialidase) gene, suggesting that this partial cluster does not confer the same function. Together, these data show that strain differences across functionally relevant gene clusters are common, indicating that statements regarding virulence of *R. gnavus* based on single isolates should be interpreted with caution. Our collection of well-characterized isolates allows researchers to assess the relevance of strain differences in future experiments.

### GWAS reveals genes related to healthy or Crohn’s-associated phenotype

In order to find genes that could explain differences in genomic repertoire of Crohn’s-derived versus healthy-derived isolates, we conducted a bacterial GWAS using Hogwash, which incorporates genomic relatedness information (Methods). On a technical note, we confirmed high correlation between core and accessory genomes (Figure 3B), and high pangenome size similarity between the Crohn’s-associated and healthy-associated groups (Figure 3C). We deemed including MAGs for this analysis to be inappropriate, as both the core and accessory genome of MAGs are substantially smaller than that of isolates (Figure S9, p < 2 × 10^−16^). Thus, their inclusion may increase false negatives or otherwise lead to spurious results.

Our bacterial GWAS analysis revealed 163 genes that were robustly associated with Crohn’s isolates (FDR < 0.05, stricter synchronous model) through a high epsilon value, which quantifies the correlation between genotype and phenotype (Figure 3D)^27^. Among the genes enriched in Crohn’s-derived isolates we found nineteen genes related to mobile genetic elements (transposases and excisionases), a predicted fucosidase which might be involved in cleaving off the terminal fucose residue on mucin, a response regulator that Bakta annotated as ‘*spo0A*’, and a holin gene (Table S3). We screened the sequence of this fucosidase gene for CAZyme domains (Methods) to gain more insight into the potential substrate preference, but found no CAZyme domains. We also compared fucosidase genes between Crohn’s and healthy isolates using CAZyme annotations for GH29 and GH95 (CAZymes with known fucose-cleaving functionality off mucin molecules), but found no significant differences (Wilcoxon rank sum test, p = 0.098 and p = 0.39, respectively; Figure S10). On the other hand, healthy-derived isolates were especially enriched for galactosidases and other genes involved in sugar metabolism (Figure 3D, Table S3). Taken together, we find novel gene-phenotype associations and provide a set of candidate genes for follow-up research on the role of *R. gnavus* in CD.

## DISCUSSION

Host phenotype-microbe association studies are often restricted to single diseases, age groups and geographic regions, which has also been the case for *R. gnavus*^10,11^. In this work we provide a detailed, global image of both the relative abundance and prevalence of *R. gnavus*, while we also investigate genomic variation within *R. gnavus* isolates in depth. In both aspects, this is to our knowledge the largest investigation to date. Key findings are the remarkably high relative abundance in newborns and young infants (Figure 1F) and the increased prevalence and abundance of *R. gnavus* in Westernized populations (Figure 1C, 1D), which have not been reported before or as comprehensively studied as here. Given its robust associations with several inflammatory diseases and allergies, many of which have high incidence in high-income countries and have their incidences rapidly increasing in newly industrialized countries^28–31^, this begs the question of whether *R. gnavus* can have detrimental immunogenic effects on the host. We show extensive genetic variation between strains in immunomodulating gene clusters, and our genetically well-characterized isolate resource can be used for experimental validation of differences in immunogenicity.

In the past decade MAGs have been increasingly used in large-scale gut bacterial genomics studies^32–36^, especially because culturing of specific gut bacteria can be highly laborious and challenging. While these MAGs have led to important biological advances, we show here that even high-quality MAGs (as defined by international standards^37^) remain of substantially worse quality than isolate genomes in multiple aspects (lower genome size and missing genes, higher GC content, amongst others, Figure 2A)^38^. In case of bacterial GWAS analyses, which aims to associate bacterial genes or genomic features with a phenotype of interest, including MAGs may therefore lead to biases and spurious associations caused by (non-)randomly missing genes due to binning and assembly artifacts. Extrachromosomal elements such as plasmids are generally not represented in MAGs, as they cannot be confidently binned, while these may be the most relevant in connection to disease and treatment options^39,40^.

Through bacterial culture combined with PacBio CCS, we have generated high-quality genome data that lead to novel insights into *R. gnavus* biology. Two aspects that highlight this are the identification of large plasmids and a conserved methylated sequence motif. To date, only one 7kb-long plasmid of *Ruminococcus gnavus* is described in GenBank (accession number NZ_CP084015.1)^41^. The two related novel plasmids we identified in the present study are much larger (164kb and 191kb; Figure S3) and likely conjugative, indicating a diversity of plasmids in *R. gnavus* that is of yet underexplored. The methylated DNA motifs that are identified here are different from those known so far (http://rebase.neb.com/cgi-bin/pacbioget?10929; Figure S4)^42^, in line with the high variability in motifs we found per genome (Figure S4). Nevertheless, we find a single m^4^C-methylated motif that is almost universally conserved across *R. gnavus* genomes (VNNVN<u>C</u>TGVNCAN). These results are reminiscent of those described for *Clostridioides difficile*^43^.

We demonstrated that *R. gnavus* is a polyphyletic species, divided into multiple (genotypically and phenotypically distinct) subspecies clades. Notably, Crohn’s-derived isolates were overrepresented in specific phylogenetic groups, while previously suggested virulence factors could not explain this separation. This suggests that these virulence factors may not play a significant role in CD symptomatology. Instead, by bacterial GWAS we identified 163 genes that could be targets for experimental validation of their role in CD development (Figure 3 and Table S3). Among these genes are 56 that we find overrepresented in CD. We listed the more noticeable candidates for which functions could be predicted. The most striking candidate is a putative fucosidase gene, as this could be directly involved in relevant cellular processes such as cell adhesion and immune system regulation^44^. Secondly, we hypothesize that genomic rearrangements and horizontal gene transfer may play an important role in the evolution of CD-associated *R. gnavus*, given the enrichment of predicted transposase and excisionase genes. Thirdly, we find a predicted holin gene which, although highly speculative, might play a role in suppressing competing bacteria^45^. A previous study identified 199 IBD-specific genes^2^, based on a pangenome of 17 draft genomes. Those draft genomes include multiple IBD-related strains and genomes from the type strain, which we find to be phylogenetically distant in our core genome phylogeny based on a pangenome of 333 genomes. This increase in genome number in the current work particularly expands the accessory genome, where the largest differences in functionality are expected. Both the previous report and our results indicate predicted functional differences in e.g. mobile elements such as transposases and (putative) mucus utilization genes underscoring the robustness of the results and narrowing down the set of target genes for IBD-specific research^2^. Furthermore, IBD research on *R. gnavus* could benefit from considering the host and possible complex host-microbe interplay for the proposed virulence factors. For example, in antibiotic-treated mice the genetic background determined whether *R. gnavus* would ameliorate or exacerbate colitis^46^.

In conclusion, we present one of the largest collections of complete genomes and associated extrachromosomal elements of any gut microbe not usually causing acute infection^47^, and provide important novel biological insight into the global epidemiology and genomic variation of *R. gnavus*. *R. gnavus* has an ambiguous relationship with human health^22^, and different strains may exert different effects on their host. Our resource of complete genomes and isolates opens promising avenues for experimental validation and further bioinformatic scrutiny, and we expect this to be valuable to the broad gut microbiome research community.

### Limitations of the study

An important consideration concerning our meta-analysis is the diversity of the studies from which the data derive. Differences in wet lab as well as dry lab methods may have led to biases in the dataset that can only be controlled for to a certain extent. Secondly, we note that although we have performed our analyses using the largest global collection of complete *R. gnavus* isolate genomes, sample size was a limiting factor for various aspects of this research. This affected our bacterial GWAS, which would benefit in terms of robustness and sensitivity especially if more and more diverse CD isolates were included. Another study limitation was the lack of experimental verification for interesting observations at the genomic level. For example, it would be of high interest to elucidate whether genomic variants of the gene cluster that is required for glucorhamnan production lead to different polysaccharides with differential immunogenicity on the host. Lastly, while this is a general limitation of (bacterial) GWAS analyses, we cannot delineate whether enriched genes in Crohn’s-derived isolates are an adaptation to an inflammatory environment or if these genes could potentially contribute to disease development and symptomatology, which would have to be assessed in preclinical models.

## MATERIALS AND METHODS

### Assessing prevalence and abundance of *R. gnavus* across human populations

We used the publicly available ‘curatedMetagenomicData’ (version 3.6.2) resource to screen 21,030 fecal metagenomes from 86 studies on all habitable continents for the prevalence and abundance of *R. gnavus*^20^. We used R (version 4.0.2; https://www.R-project.org/) to interrogate this dataset and calculate statistical parameters. We focused our analyses on metagenomes with a sequencing depth of at least five million reads and retained only the first sample per subject ID, after which 12,791 samples remained. We used the accompanying curated metadata to assess prevalence and abundance among healthy individuals across age, geography, lifestyle, and health states. Prevalence of *R. gnavus* was compared using Chi-square tests. Relative abundances were compared after adding a pseudocount of 1.3 × 10^−5^, followed by log-transformation, ANOVA, and pairwise t-tests. Sequencing depth (number of reads) was also log-transformed and compared using parametric t-test. P-values were corrected using Holm’s method and p-values ≤ 0.05 were considered significant. To compare differences in *R. gnavus* prevalence in relation to sequencing depth, we divided all Westernized and non-Westernized metagenomes in ten equal groups (quantiles) based on sequencing depth (number of reads). Relative abundances of *R. gnavus* are shown as quantiles, as adapted from previous publications^48,49^.

### Collection and curation of publicly available genome datasets

To compose a collection of *R. gnavus* metagenome-assembled genomes (MAGs) and isolate genomes, we queried a large, recent collection of gut MAGs^32^. Here, we specifically selected high-quality (HQ) MAGs annotated as *Ruminococcus gnavus* or its synonym *Faecalicatena gnavus* (with completeness > 90% and contamination < 5%)^37^. As the metadata from Almeida *et al*. does not contain curated information on disease status of the individual and this is of prime interest to our study^32^, we matched identifiers to those present in the curatedMetagenomicData package. HQ-MAGs were only included if at least both disease status and geographic origin of the original sample could be traced back. This led to a collection of 201 HQ *R. gnavus* MAGs with associated metadata.

In order to obtain additional isolate genomes to complement the MAG collection, we queried the NCBI database in December 2021 and associated metadata to retrieve at least information on disease status and geographic origin of the isolate, like the HQ-MAGs. This yielded an additional 65 *R. gnavus* isolate genomes, which all originated from China or the USA. Furthermore, we included the type strain as reference genome (ATCC 29149, accession number GCA_009831375.1)^2,26,50^.

### Metagenome-assembled genome generation from fecal metagenomes derived from multiple recurrent *Clostridioides difficile*-infected patients

We used an in-house metagenomic dataset of multiple recurrent *Clostridioides difficile*-infected patients to generate seven additional HQ *R. gnavus* MAGs – the metagenomic data of which are available in the European Nucleotide Archive under project number PRJEB44737^51^. To produce high-quality metagenome-assembled genomes (MAGs), we adapted a previously published protocol^52^.

The workflow is available as Snakemake^53^ on Zenodo (https://doi.org/10.5281/zenodo.12548294) and works as follows. Raw metagenomics sequencing reads, from which human reads had already been removed, were preprocessed using fastp (version 0.20.1, parameters: ‘--cut_right –-cut_window_size 4 –-cut_mean_quality 20 –l 75 –-detect_adapter_for_pe –y’) to trim low-quality ends, remove reads shorter than 75 bases, remove adapter sequences and remove low-complexity reads^54^. (Note: preprocessing is not part of the workflow as described on Zenodo.) Remaining, high-quality reads were assembled into scaffolds using metaSPAdes (version 3.15.4, parameters: ‘--only-assembler’)^55^. Scaffolds were binned with metaWRAP^56^ (version 1.3.2) using three binning tools: MaxBin2^57^ (version 2.2. 6), MetaBAT2^58^ (version 2.12.1) and CONCOCT^59^ (version 1.0.0) using a minimum contig length of 2500bp (‘-l’ option). Bins were then refined using metaWRAP’s ‘bin_refinement’ function, which uses CheckM^60^ (version 1.0.12) to assess bin quality, setting completeness and contamination cut-offs of 75% and 10%, respectively (‘-c’ and ‘-x’ options). After refinement, bins were reassembled using metaWRAP’s ‘reassemble_bins’ function with assemblers MEGAHIT^61^ (version 1.1.3) and metaSPAdes (version 3.13.0), again setting the minimum completeness to 75% and contamination to 10%, and the minimum length to 2000 (‘-l’ option). The resulting refined and reassembled bins were classified with the Genome Taxonomy Database toolkit (GTDB-Tk; version 2.1.0)^62^. Bins classified as *Ruminococcus gnavus* with >90% completeness and <5% contamination were included for further analyses.

### Culturing of *R. gnavus* from feces of healthy donors and patient material

We ordered *R. gnavus* strain H2_28 (DSM number 108212) from the German Collection of Microorganisms and Cell Cultures (DSMZ, Braunschweig, Germany), resuspended it in Brain Heart Infusion broth (bioMérieux, Marcy-l’Étoile, France) and streaked it on Tryptic Soy agar +5% Sheep blood (TSS; bioMérieux) to isolate pure cultures. Two unique cultures (QRD001-QRD002) were isolated from feces by streaking on Columbia Naladixic acid Agar (bioMérieux; table S1). These were all cultured in an anaerobic cabinet (Whitley A35, Don Whitley Scientific Limited, UK) with an anaerobic gas mixture (10% H_2_, 10% CO_2_, 80% N_2_) at 37°C.

To further expand our *R. gnavus* genome collection, we cultured fourteen *R. gnavus* isolates from fecal samples of healthy feces donors that were available at Vedanta Biosciences (Table S1). These were isolated and identified as follows: *R. gnavus* strains were isolated from various healthy donor stools by generating spore and non-spore fractions. Briefly, the non-spore fraction was generated by resuspending 1g of fecal material in 10mL sterile, pre-reduced PBS. The spore fraction was generated by adding 100% ethanol to the PBS fecal suspension to achieve a 50% (v/v) ethanol concentration. The fecal ethanol suspension was incubated at 25°C for 1hr while shaking. Following incubation, the fecal ethanol suspension was centrifuged at 3400×g for 20 minutes and the cell pellet resuspended in 1mL of sterile, reduced PBS. Serial dilutions of the spore and non-spore fraction were plated on either Eggerth-Gagnon + 5% horse blood agar, Brucella Blood Agar (Anaerobe Systems, Inc., Morgan Hill, California, USA), MSAT (Anaerobe Systems), or chocolate agar and incubated at 37°C anaerobically for 72hr. Isolated colonies were identified by Sanger sequencing of the 16S amplicon using 8F and 1492R primers and Illumina shotgun sequencing. Isolated colonies were inoculated into 1.2mL of Peptone Yeast Extract Broth with Glucose (PYG; Anaerobe Systems) in a 96-deep well plate and incubated at 37°C anaerobically for 48hr. After incubation, colony identity was determined by performing PCR from 200µL of the culture using universal 16S primers 8F and 1492R. Selected isolates were then sub-cultured from the 96-deep well plate onto the appropriate agar medium and incubated at 37°C anaerobically for 72hr. An isolated colony from this plate was inoculated into 5mL of PYG and incubated at 37°C anaerobically for 24hr. 1mL of the culture was pelleted by centrifuging at 10000×g for 5 minutes. DNA was extracted from the pellet using the DNeasy blood and tissue kit (Qiagen, Hilden, Germany) following the manufacturer instructions. Colony identity was determined again by Sanger sequencing of the 16S gene amplicon using 8F and 1492R primers and Illumina shotgun sequencing.

Furthermore, fourteen isolates were cultured and collected at the University Medical Center Groningen as follows. Brucella blood agar medium (Mediaproducts BV, Groningen, The Netherlands) was used to cultivate the *R. gnavus* strains QRD024, QRD025 and QRD028 from human clinical specimens (Table S1). The plates were transferred to an anaerobic workstation (Whitley A45) after inoculation and incubated for one to three days at 37°C. The anaerobic medium YCFA supplemented with either apple pectin or porcine mucin type III (4.5 g/l) was used for the isolation of QRD026, QRD027, and QRD029-QRD031 as described earlier^63^. Fecal samples of healthy volunteers were used for inoculation on pre-reduced medium and the plates were incubated at 37°C in an anaerobic chamber (Whitley A35 Workstation) with an anaerobic gas mixture (10% H_2_, 10% CO_2_, 80% N_2_). The strains QRD032-QRD037 were isolated from fecal samples of IBD patients on either phenylethyl alcohol agar (Mediaproducts BV, Groningen, The Netherlands), brain heart infusion agar (Oxoid Limited, Cheshire, UK) supplemented with yeast (2,5 g/l), hemin (0,001% w/v) and cysteine (1 g/l) or YCFA medium supplemented with glucose (4.5 g/l).

Moreover, isolates as cultured in their respective publications were obtained from the Broad Institute^13^, and Sanger Institute^64^. All cultures from outside the Leiden University Medical Center (LUMC) were sent to the LUMC as frozen glycerol stocks and anaerobically cultured on TSS. After obtaining pure colonies, all isolates were independently confirmed to be *R. gnavus* in our laboratory using matrix-assisted laser desorption/ionization coupled to a time-of-flight mass spectrometer (MALDI-TOF; Bruker Daltonics GmbH, Bremen, Germany). All isolates were able to grow on TSS, CNA and Chocolate agar PolyViteX (bioMérieux) and the colony morphology appeared on plates as round, glassy white colonies with a bright white center. Sometimes colonies displayed concentric circles, reminiscent of checker game pieces.

### Data processing of Illumina-sequenced *R. gnavus* isolates

The fourteen isolates cultured at Vedanta Biosciences were sequenced on the Illumina NextSeq platform using 150bp paired-end reads. These data were included with the isolate short-read-based genomes, increasing the number to 79 short-read isolates. Raw Illumina sequence data was cleaned and trimmed using fastp (v0.23.2) and sequence quality was inspected using Fastqc^65^ (v0.11.9) and Multiqc^66^ (v1.8). Cleaned reads were assembled by first using SKESA^67^ (v2.4.0) and subsequently SPAdes (v3.15.3) with “--untrusted-contigs” and “--isolate” parameters.

### Quality control and annotation of short-read-based genome collection

We have collected a total of 287 short-read-based genomes of *R. gnavus*, consisting of 79 assembled whole-genome sequences from cultured isolates and 208 metagenome-assembled genomes (MAGs). We also added the one available reference sequence in our analyses (NCBI GenBank accession number GCF_009831375.1). We filtered out contigs shorter than 1,000 bp using BBtools’ reformat.sh (version 37.62; https://sourceforge.net/projects/bbmap/). We estimated completeness and contamination of all genomes using CheckM (version 1.0.13) and verified that all genomes taxonomically classify as *R. gnavus* using GTDB-Tk (version 2.1.0). Assembly length statistics were determined using QUAST^68^ (version 5.0.2). Finally, genomes were annotated using Bakta^69^ (version 1.6.1), which also provides the number of open reading frames, or predicted genes, per genome.

### DNA isolation of *R. gnavus* isolates and generation of complete genomes using PacBio circular consensus sequencing

To generate complete genomes, 45 isolates were subjected to long read sequencing on the Pacific Biosciences (PacBio, Menlo Park, California, USA) Sequel IIe platform at the Leiden Genome Technology Center. To prepare high molecular weight total DNA, isolates were cultured anaerobically overnight in 10 mL BHI at 37°C. Cells from 5 mL of culture were pelleted and processed using the Qiagen Genomic-tip 100/G, according to the manufacturer’s instructions. SMRTbell® libraries were generated as follows. Genomic DNA was sheared with the Megaruptor 3 system (Diagenode LLC, Denville, New Jersey, USA) using 35 cycles. Libraries were generated according to the following manufacturer’s procedure and checklist: Preparing whole genome and metagenome libraries using SMRTbell® prep kit 3.0 (PN 102-166-600 REV02 MAR2023), thereby using barcoded adapters. Size-selection was performed on library sub-pools using either diluted AMPure PBbeads (PacBio, 35% beads, 3.1x v/v ratio) or Blue Pippin (Sage Science, Beverly, Massachusetts, USA), depending on the insert-size of the libraries. The libraries were sequenced on a PacBio Sequel IIe platform with a 30 hour movie time using Sequel II Binding Kit 3.2 and Sequel II sequencing kit 2.0.

Raw PacBio reads were assembled using five different assemblers: Canu^70,71^ (version 2.2), Flye^72^ (version 2.9.2), Raven^73^ (version 1.8.1), Hifiasm^74^ (version 0.19.6-r595) and IPA (version 1.8.0; https://github.com/PacificBiosciences/pbipa). Note that sample QRD034 was sequenced much deeper than the rest and subsampled to 30% of reads (= 277X coverage) to facilitate assembly. Contigs were taxonomically classified using the Contig Annotation Tool (CAT version 5.2.3)^75^ to verify if they derived from *R. gnavus*. Canu, Flye, Hifiasm and IPA report if assembled contigs are linear or circular. From the different assemblies, we selected the assembly that yielded the longest contig and the longest total assembly length (all exceeding 3 Mb), giving Flye precedence as it provides the most extensive statistics. All contigs from selected assemblies were reoriented using dnaapler^76^ (version 0.3.0) to start at the *dnaA*, *repA* or *terL* gene for chromosomes, plasmids and bacteriophages, respectively. Raven and Hifiasm produce assembly graphs, which were viewed to assess if contigs were linear or circular. Assemblies with a smaller secondary circular contig were analyzed with geNomad^77^ (version 1.7.4) to predict the probability of it being a plasmid, using the built-in score calibration module with aggregated results from both the marker-based and neural net-based classifications.

We included two isolates derived from the strain DSMZ 108212, of which one we obtained directly from the DSMZ (QRD005) and the other was cultured at the Sanger Institute (QRD022). Assembly with Hifiasm yielded a 3.3Mb contig and a 28kb contig for QRD022, while QRD005 could not be resolved to less than three contigs, with the longest being 2.4Mbp. These two assemblies were not completely identical and we decided to use a reference-based assembly of the unresolved one against the 3.3Mb contig using minimap2^78^ (version 2.29) and samtools^79^ consensus (version 1.19; parameters: ‘--min-MQ 5 –-min-depth 10’) to generate an improved assembly of QRD005. This resulted in two contigs of 3.3Mb and 178bp. We manually removed the 178bp fragment and use the single 3.3Mb contig assembly as representative of the ‘DSMZ-108212’ = QRD005 isolate (Table S1).

Final genome assemblies were annotated with DNA methylation information from the PacBio SMRT Link Microbial Genome Analysis platform.

### Antibiotic resistance screening of isolate genomes

To assess the genotypic antibiotic resistances in isolate genomes, we screened 79 short-read genome sequences of isolates, the 45 newly generated long-read genomes, and the one reference genome for the presence of antibiotic resistance genes using ABRicate (version 0.8.13; https://www.github.com/tseemann/abricate) with NCBI’s AMRFinderPlus database (downloaded 11 November 2022, containing 5,735 sequences)^80^. Genes were assumed present if at least 95% of the gene matched with at least 95% identity to the gene in the database. For *in vitro* validation, ten isolates – five with *tet* tetracycline resistance genes and five without – were assessed for tetracycline minimum inhibitory concentrations (MIC at 48h) using an ETEST (bioMérieux) on TSS medium at 37°C in a Whitley A35 anaerobic cabinet.

### Search for previously described inflammatory factors of *R. gnavus*

Several *R. gnavus* genes have previously been associated with intestinal inflammation. We screened our collection of genomes for the presence of two superantigen genes (accession numbers WP_105084811.1 and WP_105084812.1)^15^, 23 genes encoding the machinery to produce a proinflammatory (glucarhamnan) polysaccharide (NZ_AAYG02000032.1)^13^, one tryptophane decarboxylase gene (RUMGNA_01526 from UniProt)^16^, and 20 genes encoding a capsule polysaccharide (RUMGNA_02411 – RUMGNA_02392 from UniProt)^14^. We used protein BLAST^81^ (blastp; version 2.13.0) to screen the genomes for the presence of each of these genes. Only hits that covered at least half of the gene of interest (‘-qcov_hsp_perc 50’) with an E-value of 1 × 10^−20^ or smaller (‘-evalue 1e-20’) were considered for further analysis. Gene clusters were considered present when all the genes were detected.

Using the same method, we also screened genomes for the presence of the bilirubin reductase gene (*bilR,* WP_009244284.1)^82^, selenium-dependent xanthine dehydrogenase (*sd-XDH*, QHB24869.1)^17^, and the *nan* cluster for sialic acid metabolism (RUMGNA_02691 through RUMGNA_02701 from UniProt)^18^.

### Annotation of functional pathway genes

We annotated carbohydrate-active enzymes (CAZymes) by comparing the genomes to dbCAN^83^ (version 10) using HMMer^84^ (version 3.3.2). Within the CAZyme families, we focused on two glycosyl hydrolase families that include fucosidases, GH29 and GH95, which have been described as important for mucus utilization^85^, a main feature of *R. gnavus*. Genomes were also annotated using KEGG-Decoder^86^. Pathways for chemotaxis and flagellum biosynthesis were annotated using the KOALA definitions available online^23^. Moreover, genomes were screened for the presence of annotated biosynthetic gene clusters (BGC) using antiSMASH^87^ (version 6.1.1).

### Comparison of whole genomes to find clusters of genomic variants

Whole genomes were compared to one another using average nucleotide identity (ANI) with fastANI (version 1.33)^25^. Furthermore, genomes were subjected to a pangenome analysis using Panaroo (version 1.3.0; parameters ‘--clean-mode strict –a core –-aligner mafft –-core_threshold 0.95’)^88^. For the pangenome, we considered genes that occur in at least 95% of genomes core genes as recommended when including MAGs^89^. The core genes were concatenated and using MAFFT^90^ (version 7.505) a core genome multiple sequence alignment was generated, which was automatically trimmed using trimAl^91^ (version 1.4.1). A maximum likelihood phylogeny was inferred from the trimmed multiple alignment using IQ-tree^92^ (version 2.2.0.3), including ModelFinder Plus^93^ to automatically select the best fitting evolutionary model and ultrafast bootstrap (1000 replicates) to calculate branch support^94^. The selected models were: short-read genomes GTR+F+I+I+R9; long-read genomes GTR+F+R7; all genomes GTR+F+R10. A Principal Coordinate Analysis (PCoA) of core genome phylogenetic distances was conducted using the function ‘cmdscale’ from the stats R package, followed by k-means clustering using ‘kmeans,’ also from the stats package.

### Bacterial genome-wide association study (GWAS)

To identify genes that are putatively associated with CD, we subjected genomes of *R. gnavus* isolates to a bacterial genome-wide association study using Hogwash (version 1.2.6; parameters: ‘fdr = 0.05, bootstrap = 0.875, grouping_method = “post-ar”’)^27^. Genomes were assigned healthy or CD phenotype based on available metadata on health status from the person from whom the *R. gnavus* isolate was cultured. We included short-read sequencing isolate draft genomes as well as our in-house generated PacBio complete genomes. If multiple sequences of the same isolate existed, we deduplicated based on ANI > 99.9%. Of these duplicates, we picked the first based on alphabetic order as representative, and we preferentially select long-read-based genomes when available.

Hogwash reconstructs the evolutionary history of the genomes of interest using a phylogenetic tree and predicts where genotype and phenotype transitions occurred to assess where genotype and phenotype transitions coincide. We made use of the high correlation between core and accessory genome to use these two as input, together with phenotype of either CD or healthy. This resulted in fourteen *R. gnavus* isolate genomes derived from CD patients and 41 from healthy people (total N = 55). We used a matrix of (accessory) gene presence and absence generated by Panaroo as input for Hogwash. As phylogenetic tree, we pruned the tree of all *R. gnavus* genomes inferred by IQ-tree to include only this set of 55 deduplicated genomes and midpoint rooted it.

### Statistical analyses

All tools were run with default parameters unless stated otherwise. Statistical analyses and visualization were done in R (version 4.0.2) using RStudio (https://posit.co/). A p-value of 0.05 or smaller was considered significant.

## DATA AND CODE AVAILABILITY

The long-read whole-genome sequencing data and corresponding assemblies of isolates presented in this study are available from the European Nucleotide Archive under accession number PRJEB76407. A complete set of short and long-read genomes together with metadata is available through Zenodo (https://doi.org/10.5281/zenodo.12548497). Scripts of both the whole-genome annotation and comparative genomics analyses, as well as further downstream and statistical analyses are available on Zenodo (https://doi.org/10.5281/zenodo.12548546).

## ACKNOWLEDGEMENTS

This study was supported by the Leiden University Fund / Dr. F.F. Hofman Fonds, (www.luf.nl) to QD. Further funding was provided by the LUMC (LUMC Fellowship to G.Z.), the Health + Life Science Alliance Heidelberg Mannheim through state funds approved by the State Parliament of Baden-Württemberg (postdoctoral fellowships to Q.D. and N.K.) and EMBO postdoctoral fellowship (ALTF 1030-2022 to Q.D.). The Graduate School of the Medical Sciences of the University of Groningen provided a grant to N.P. We thank all members of the scientific community that generously provided us with *R. gnavus* isolates.

## AUTHOR CONRTIBUTIONS

Conceptualization, EK, WKS and QD; methodology, NP, IS, LS, AvdM, ET, NK, RV, SK, HH, KF; investigation, SN, NP, RV, IS, LS and QD; formal analysis, SN and QD; writing – original draft, SN and QD; writing – review & editing, all authors; supervision, SK, GZ, EK, WKS and QD; funding acquisition: QD

---

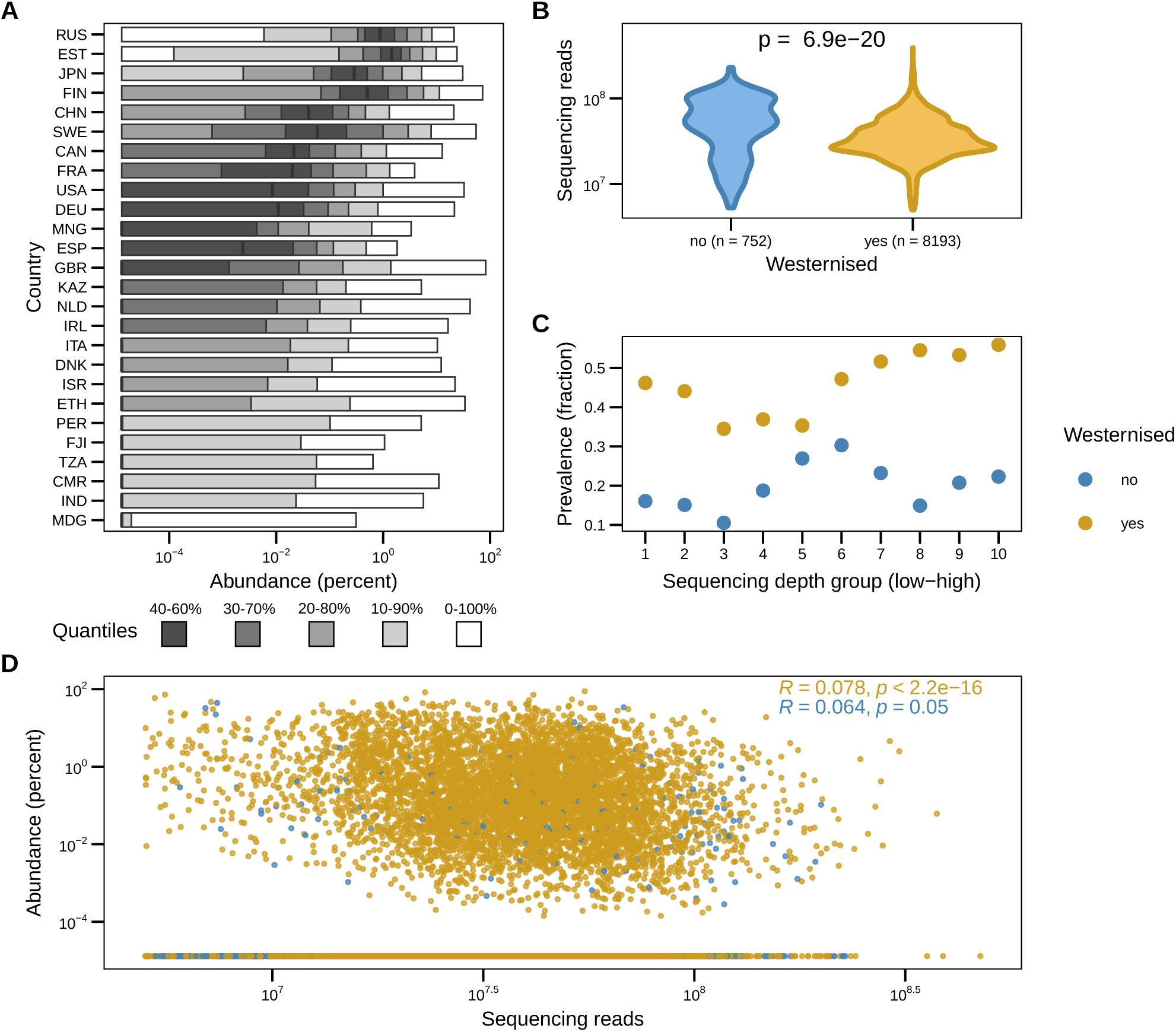

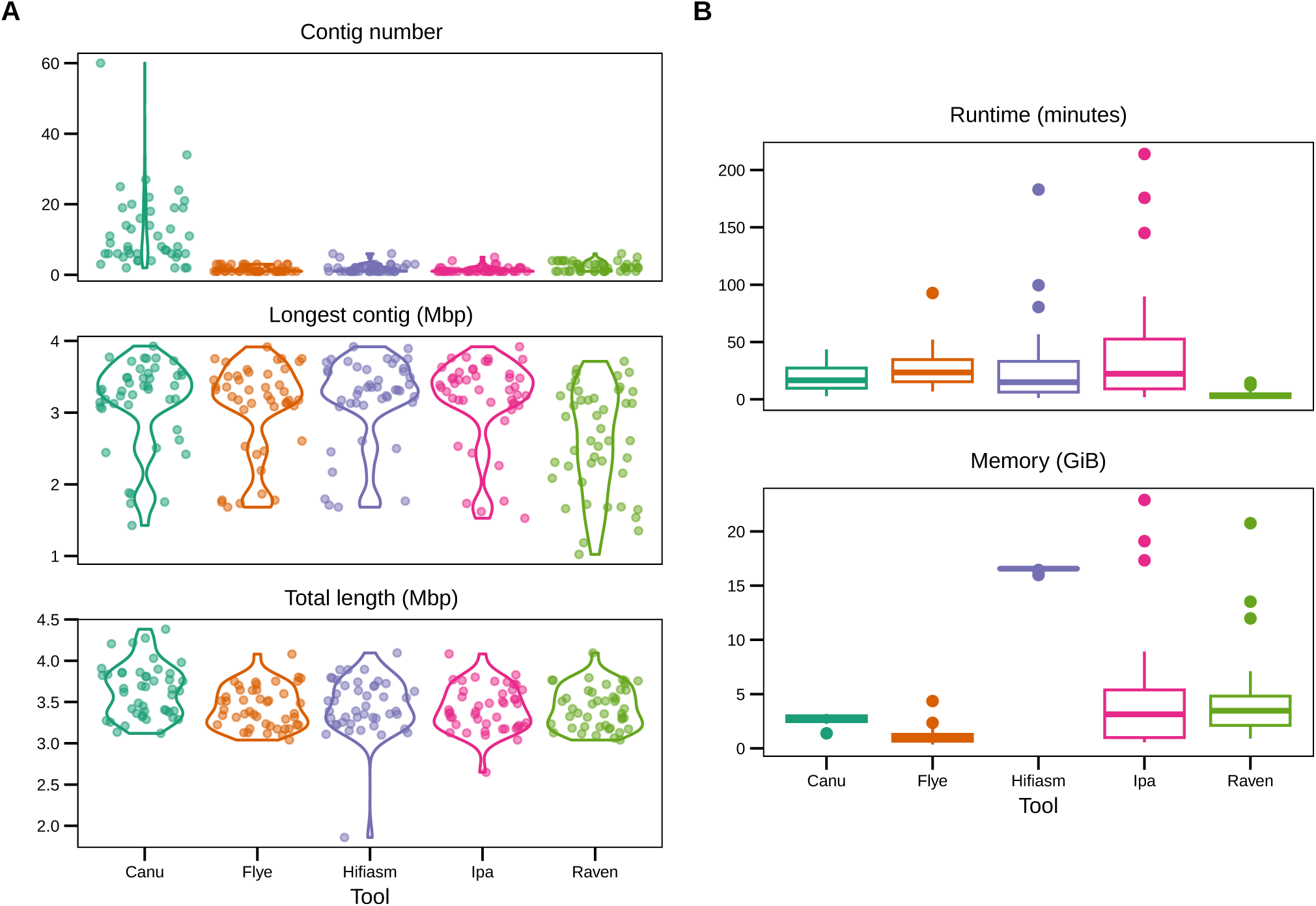

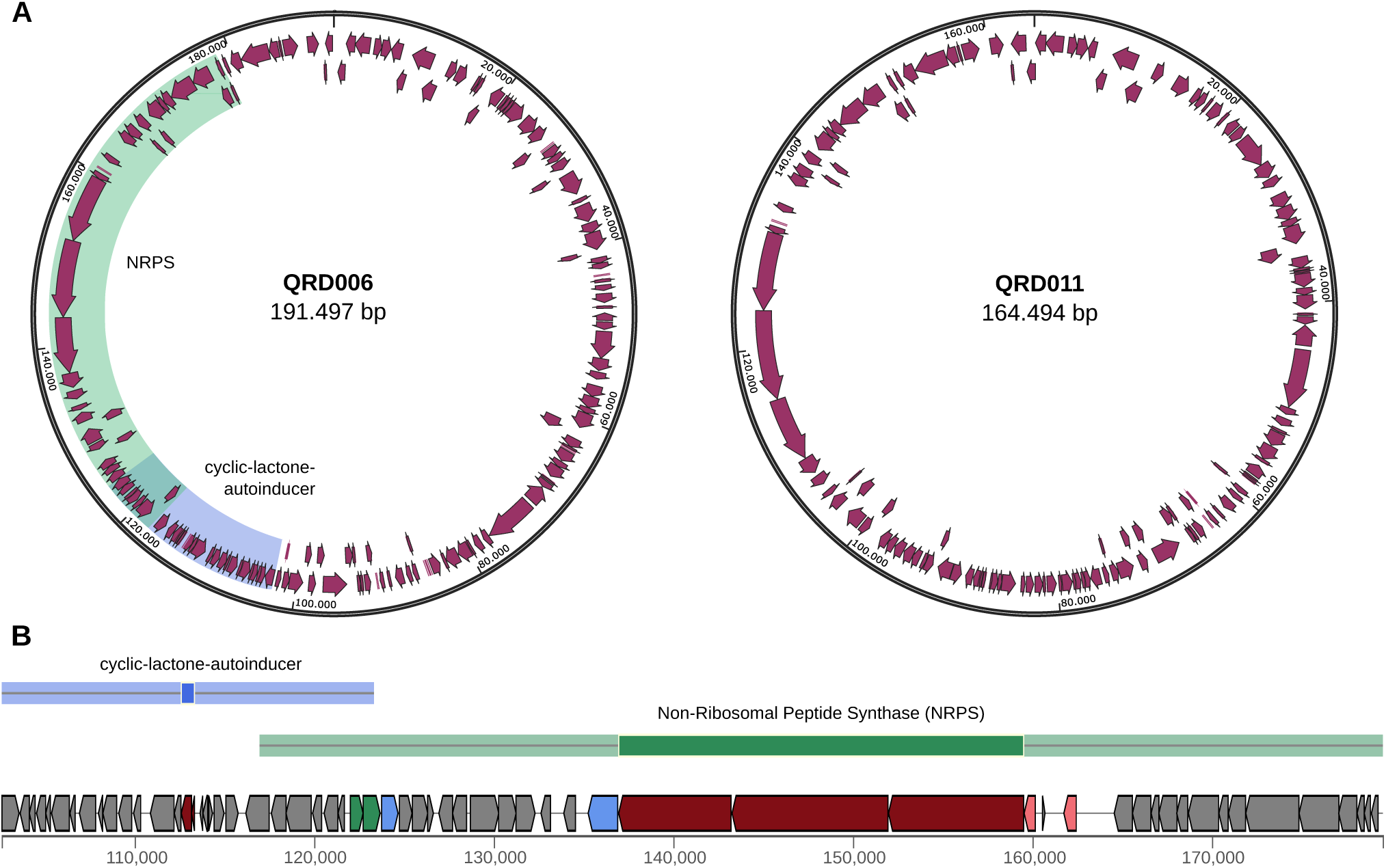

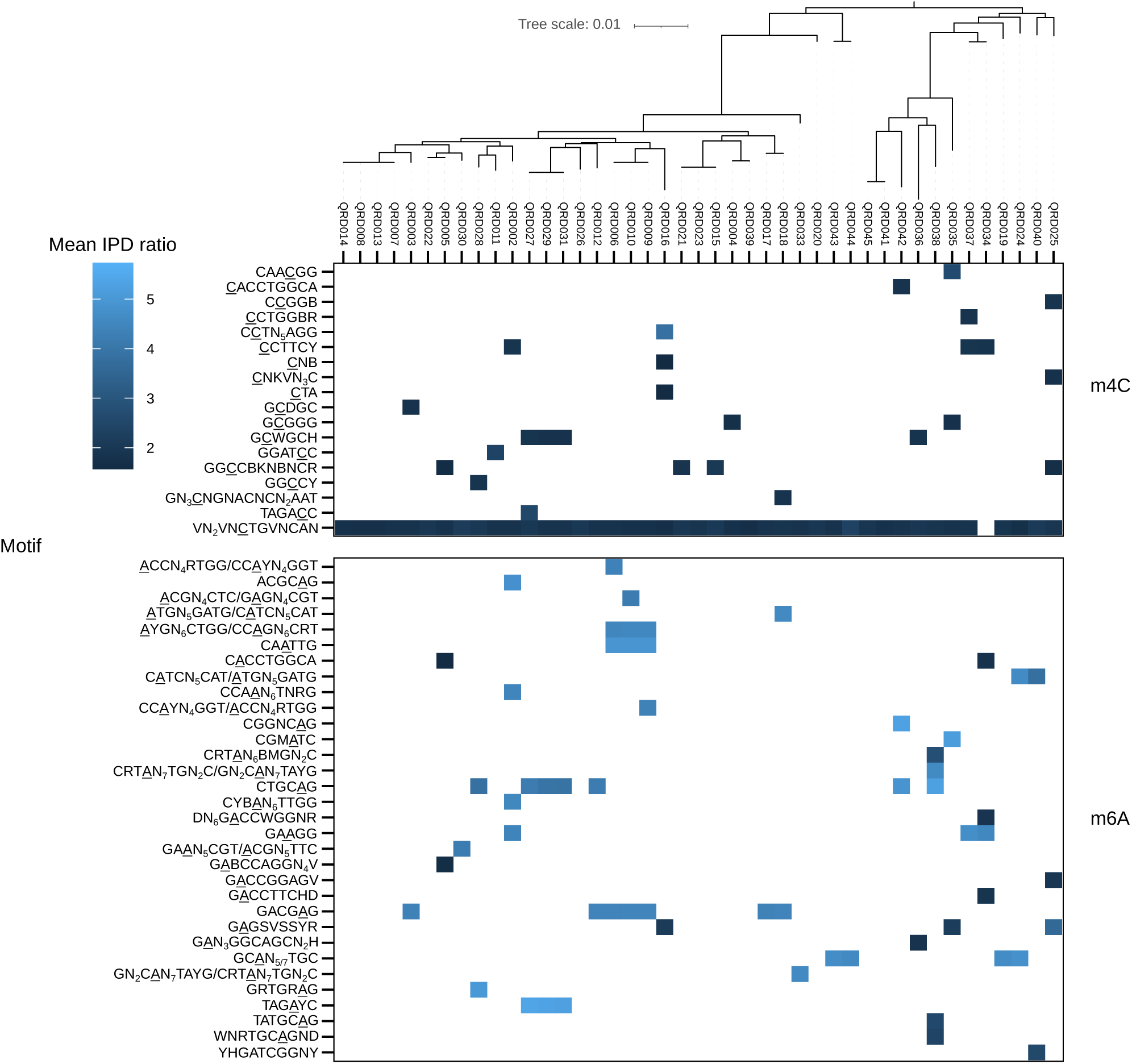

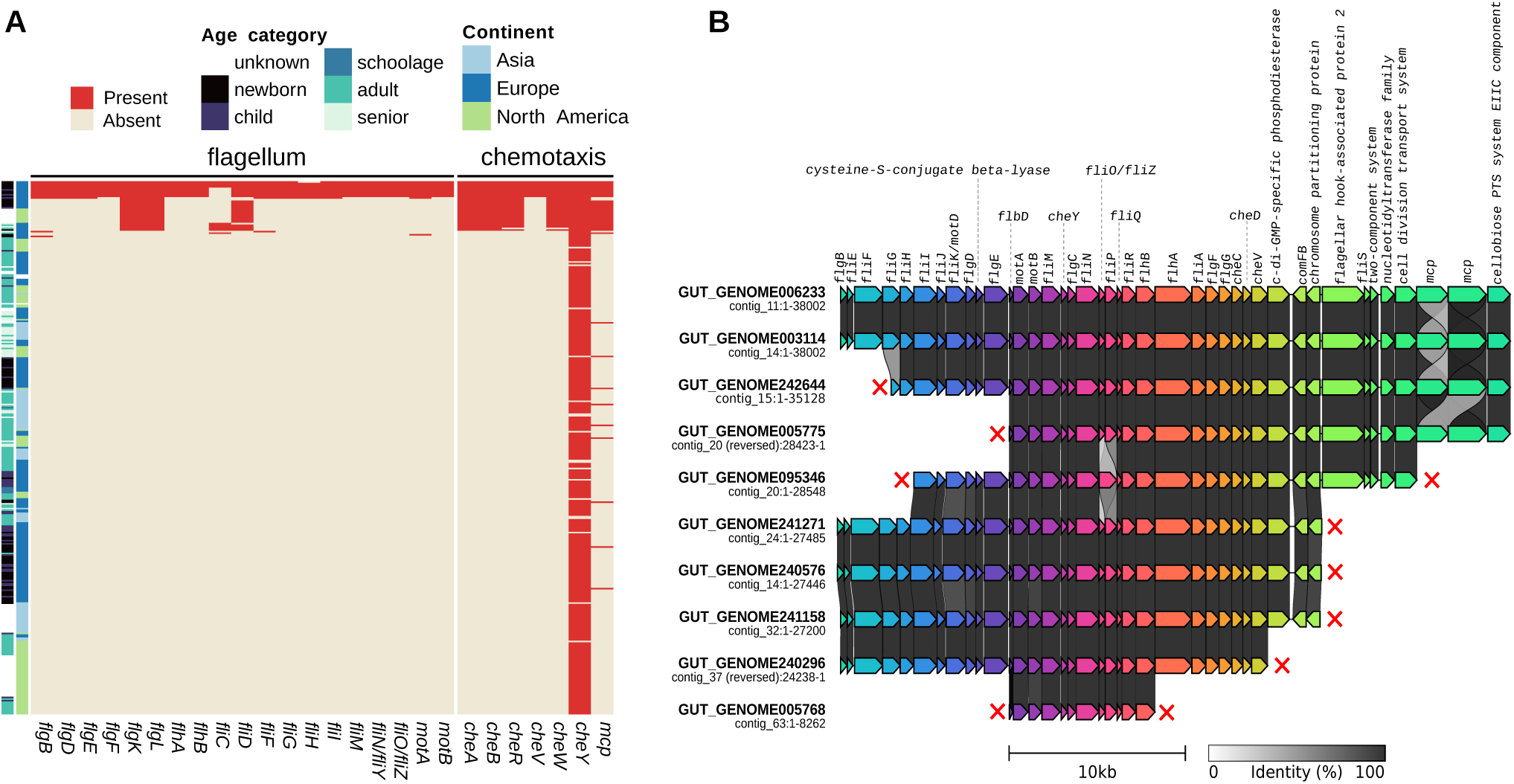

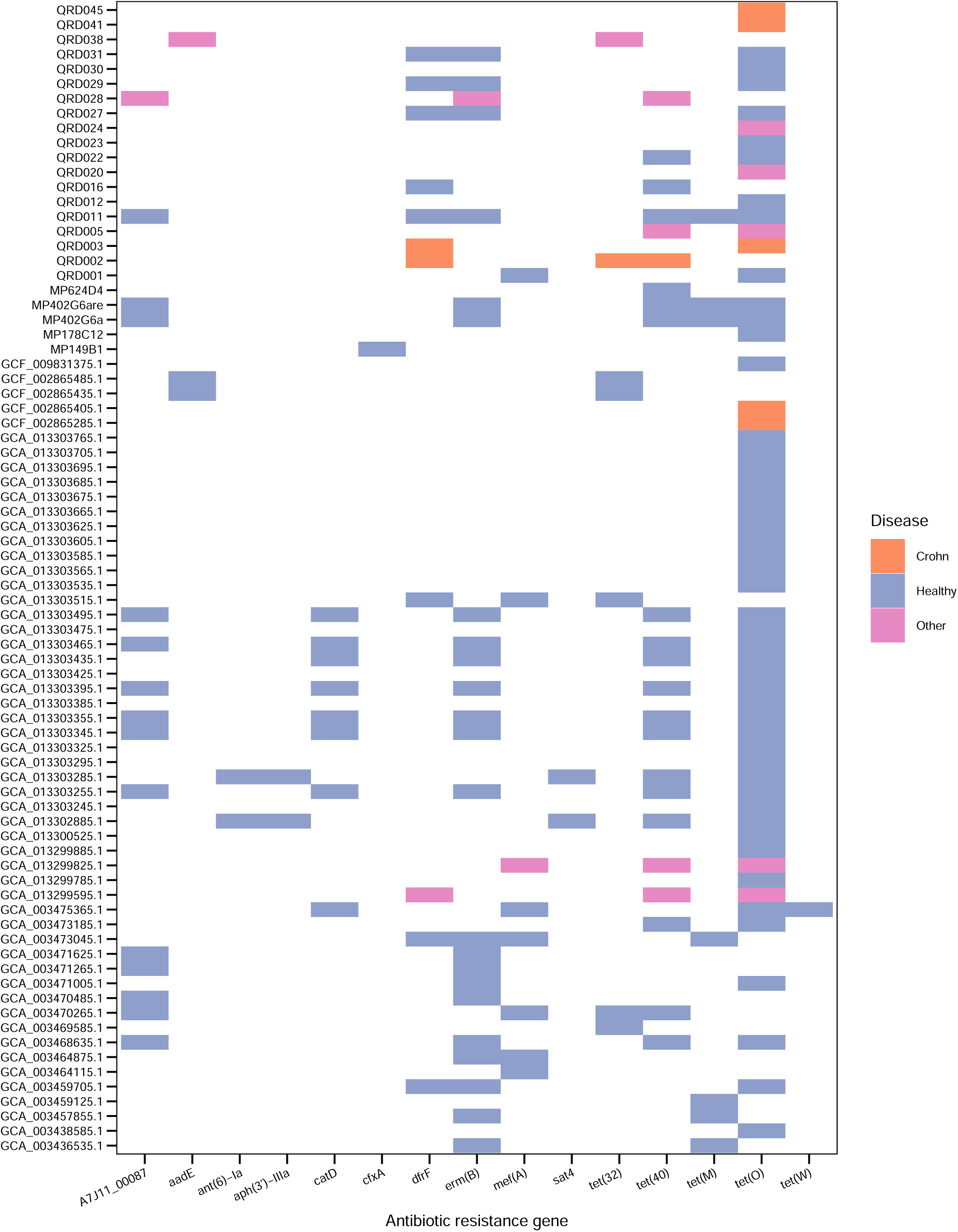

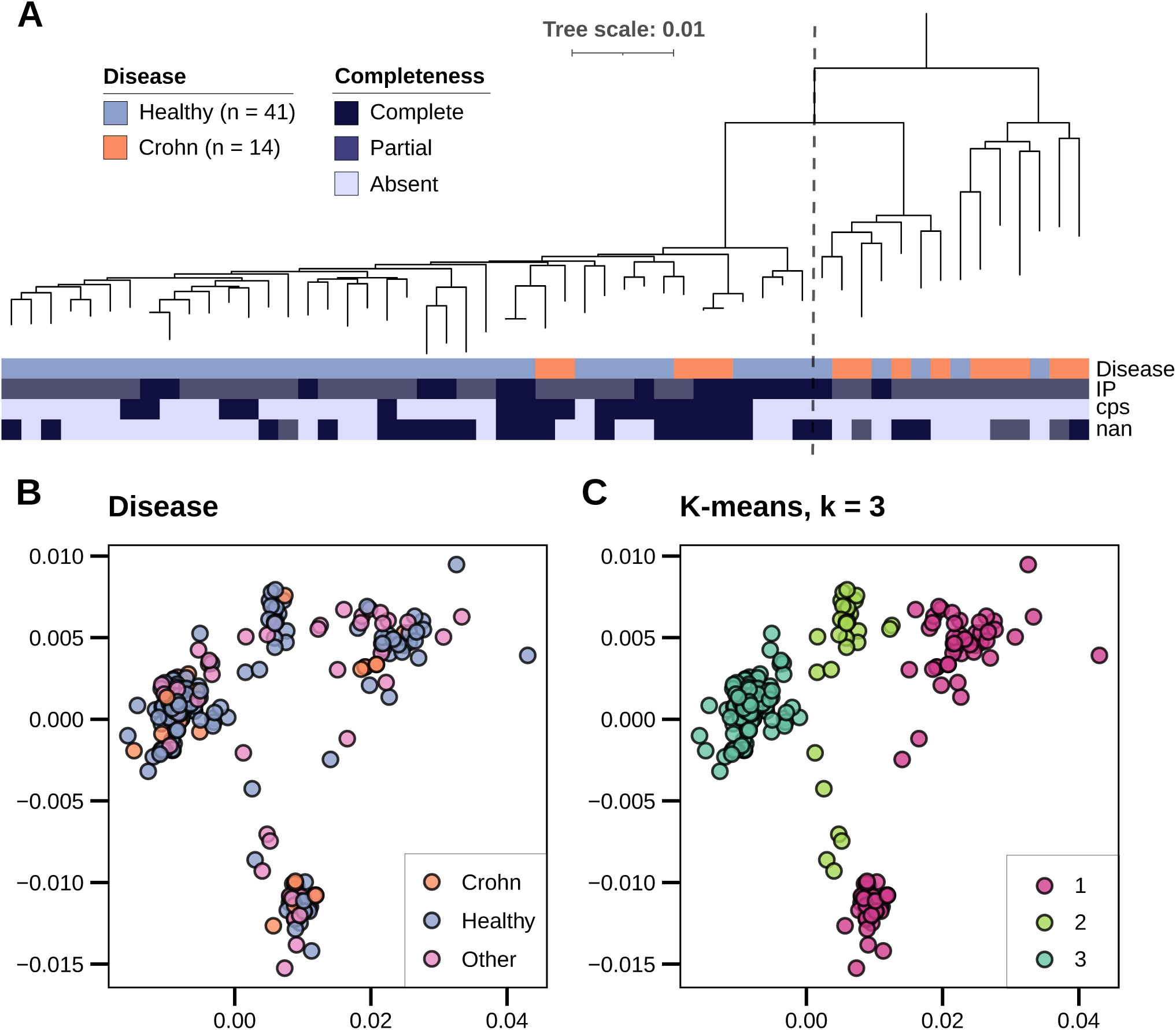

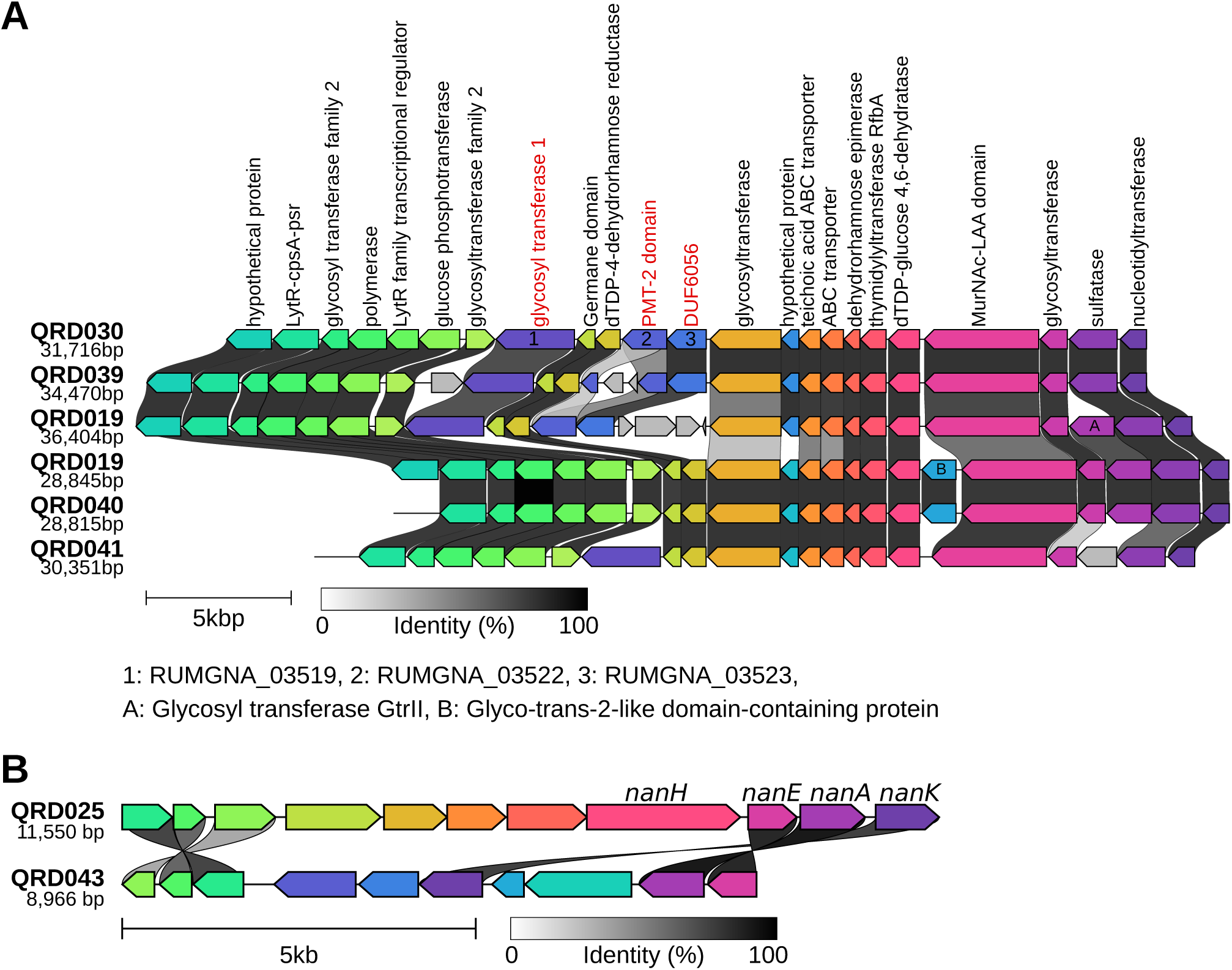

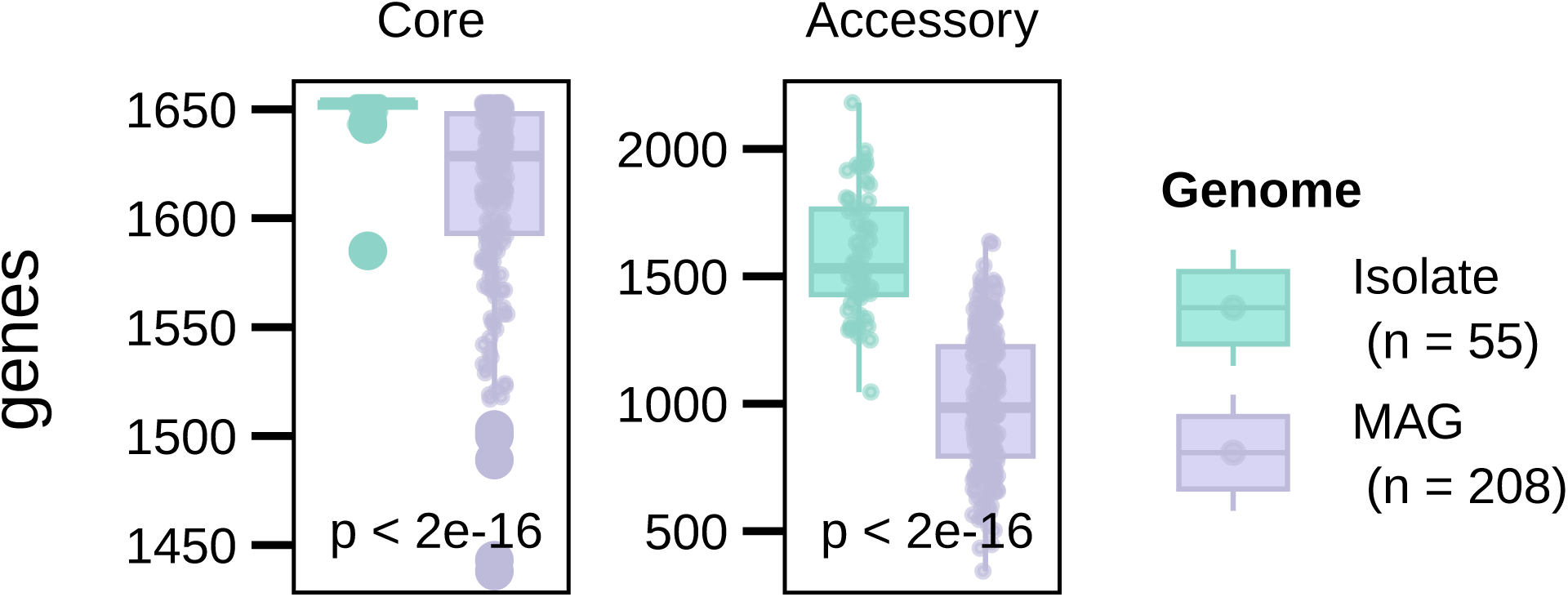

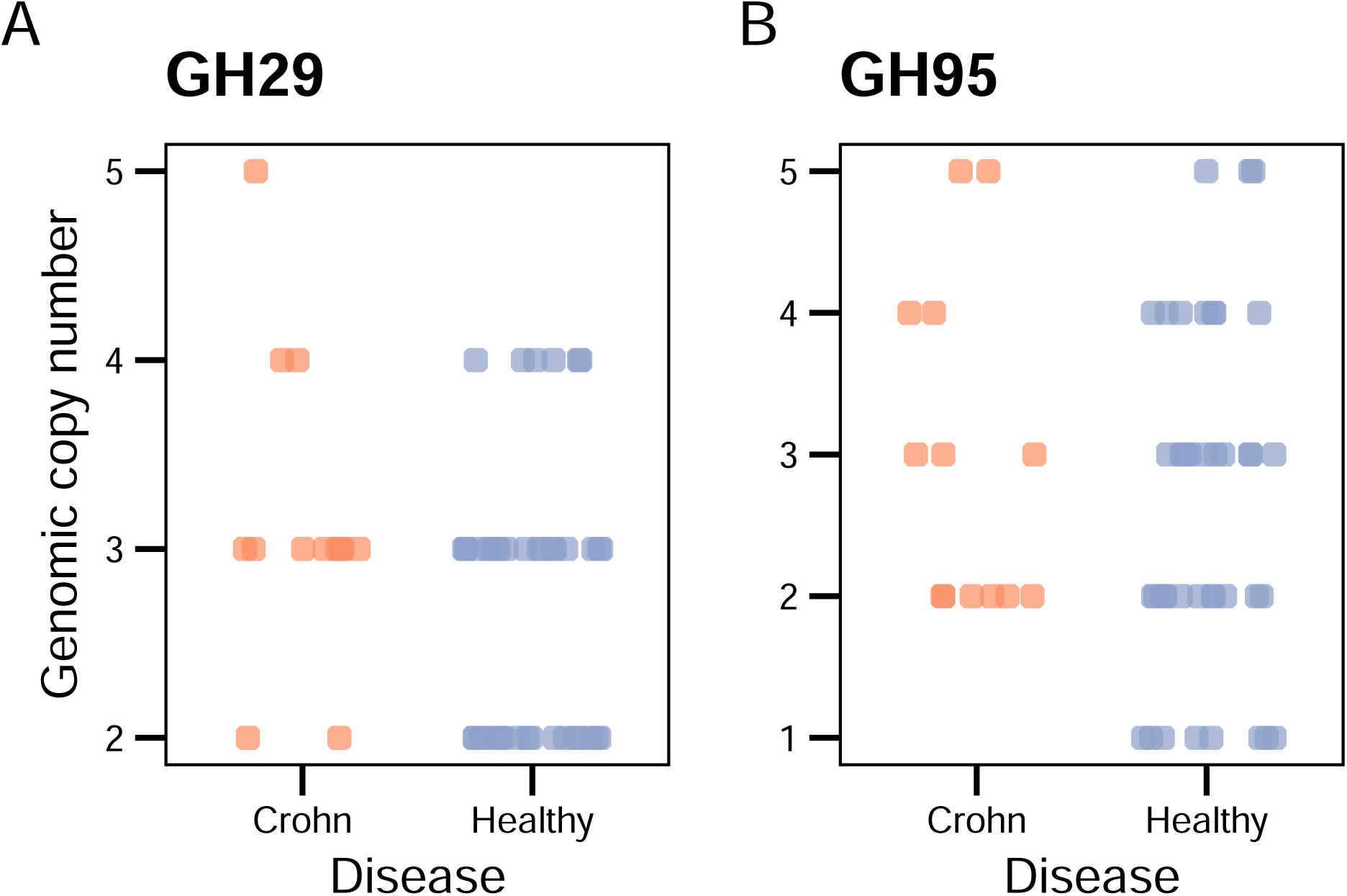

