## Supplemental table S2 and supplemental figures S1-S10 for "Metagenomic global survey and in-depth genomic analyses of *Ruminococcus gnavus* reveal differences across host lifestyle and health status"

**SUPPLEMENTARY TABLES AND FIGURES**

**Table S2. Antibiotic resistance of tested isolates, related to figure 2.**

| **Isolate** | **TET_genes** | **tetracycline_MIC** |
| --- | --- | --- |
| QRD033 | - | 0.047 |
| QRD035 | - | 0.064 |
| QRD004 | - | 0.064 |
| QRD039 | - | 0.064 |
| QRD013 | - | 0.064 |
| QRD027 | tet(O) | 2 |
| QRD030 | tet(O) | 8 |
| QRD016 | tet(40) | 16 |
| QRD022 | tet(O), tet(40) | 16 |
| QRD002 | tet(40) | 48 |

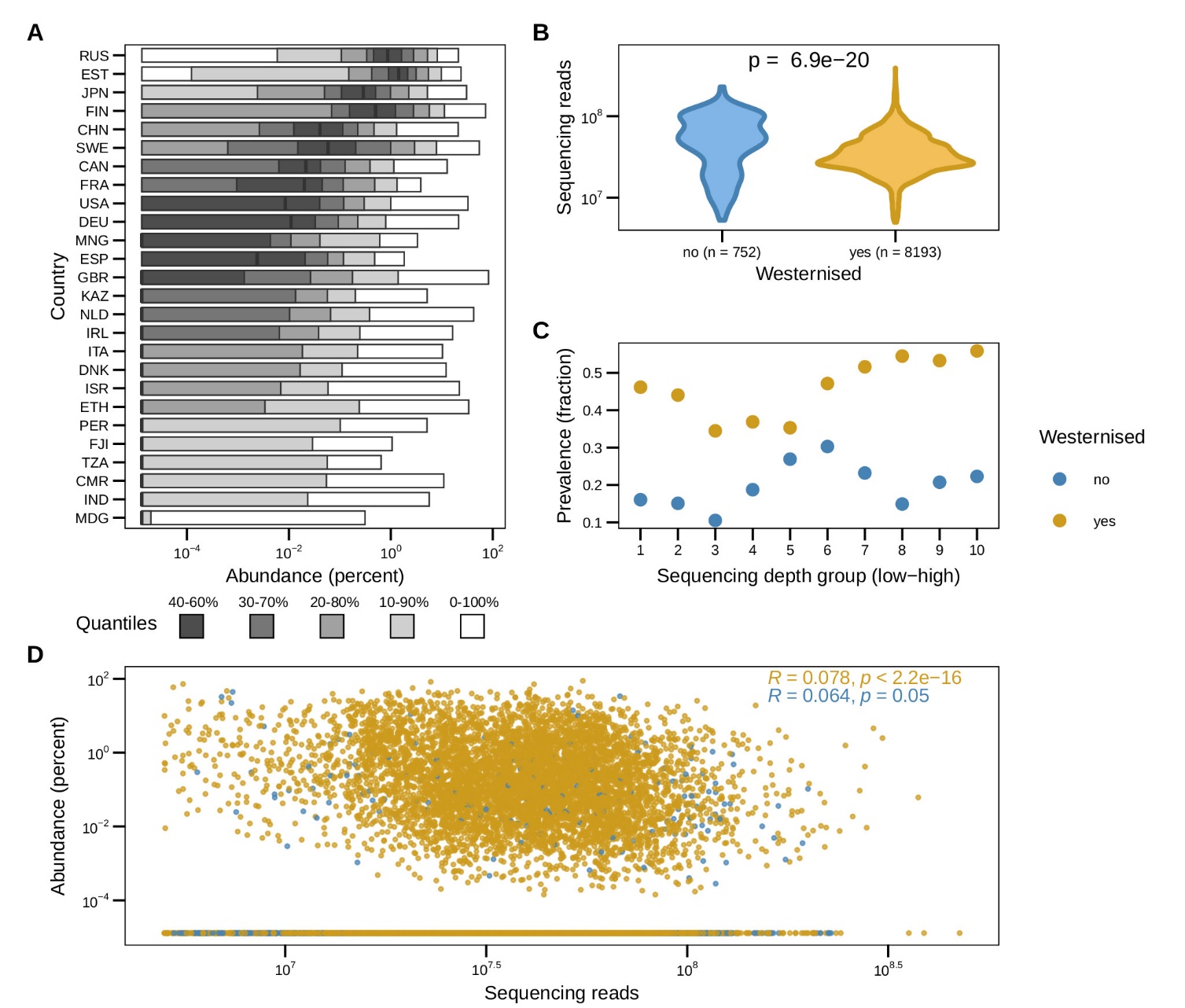
**Figure S1. Abundance of *Ruminococcus gnavus* per country and sequencing depth controls for Westernised versus non-Westernised, related to Figure 1.**

(A) Relative abundances of *R. gnavus* per country.

(B) Comparison of sequencing depth between samples collected from Westernised and non-Westernised people. We applied the student’s t-test to evaluate the difference in sequencing depth in number of reads per sample.

(C) Comparison of prevalence of *R. gnavus* between Westernised and non-Westernised samples divided in 10 equal quantiles.

(D) Comparison between sequencing depth and relative abundance of *R. gnavus*, separated by Westernisation status. Correlations are calculated using Spearman’s correlation statistic.

A pseudocount of 1.3*10^-5^ is added to all abundances to enable visualisation on a logarithmic scale (panels A, & D).

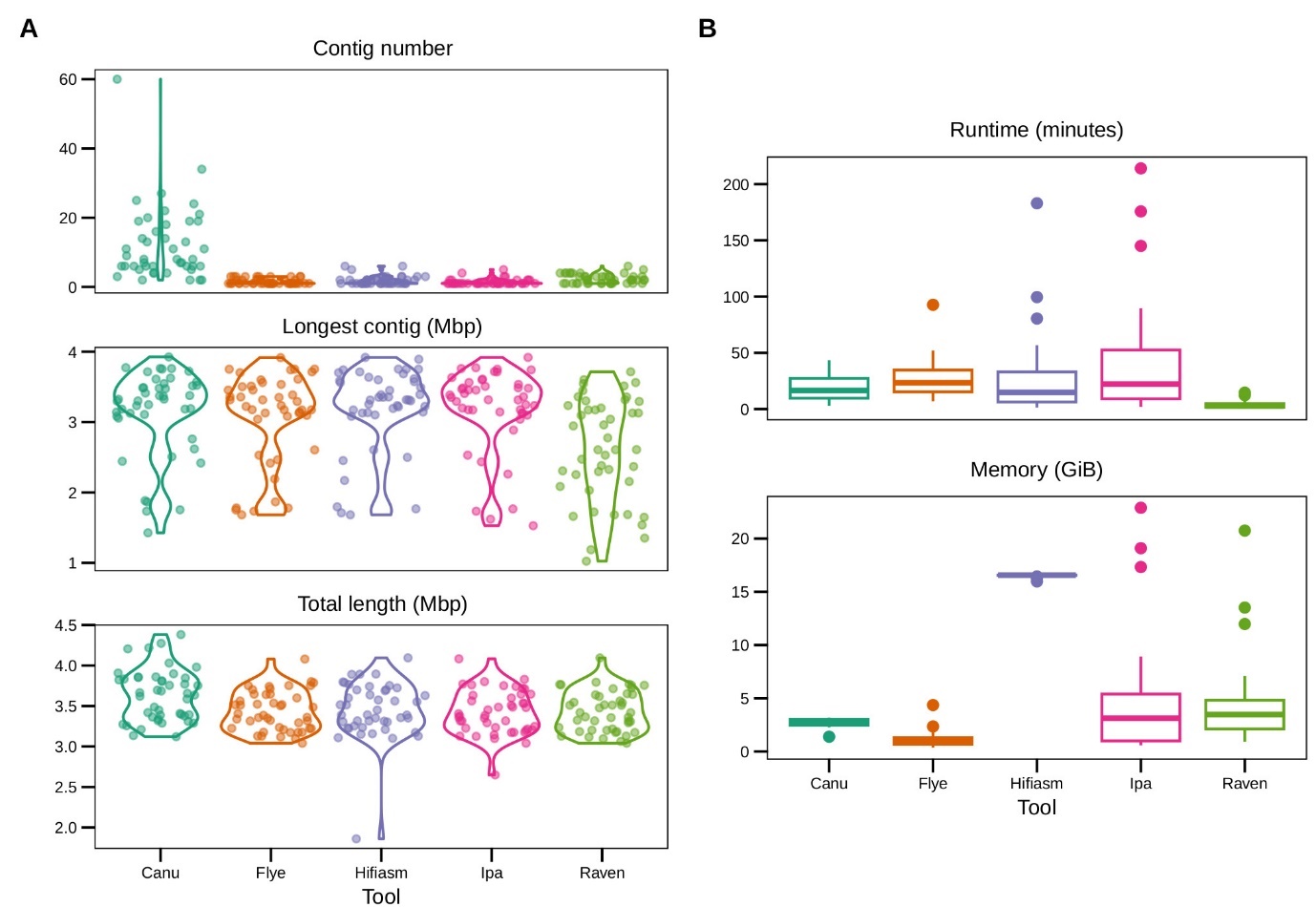

**Figure S2. Benchmark of *de novo* assembly methods for bacterial long-read sequencing data related to Figure 2.**

(A-B) We tested five different assemblers (Canu, Flye, Hifiasm, IPA, and Raven) on our 45 samples of PacBio HiFi reads of *Ruminococcus gnavus*. This figure shows the median result of three replicates.

(A) Number of contigs, the length of the longest contig and the total length of contigs generated by the assembly tool.

(B) Runtime and memory use of each assembly tool.

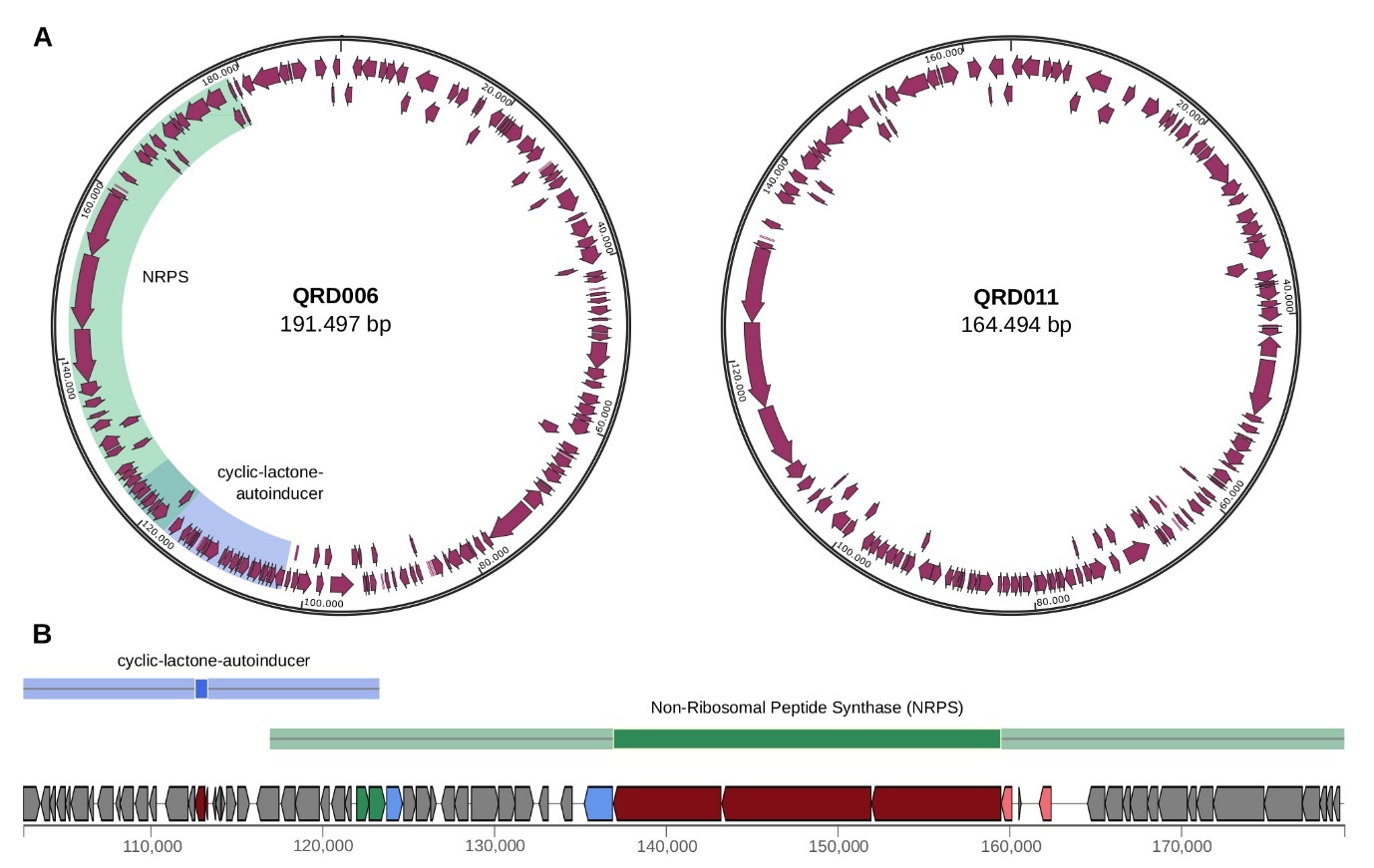

**Figure S3. Two novel large plasmids with a biosynthetic gene cluster, related to figure 2.**

(A) Circular representation of two plasmids identified in isolates QRD006 and QRD011, with predicted open reading frames shown as arrows.

(B) Biosynthetic gene cluster as identified on the plasmid from isolate QRD006. Numbers on the x-axis represent genomic locations in basepairs. Positions of the predicted gene cluster are also highlighted in (A) in corresponding colors.

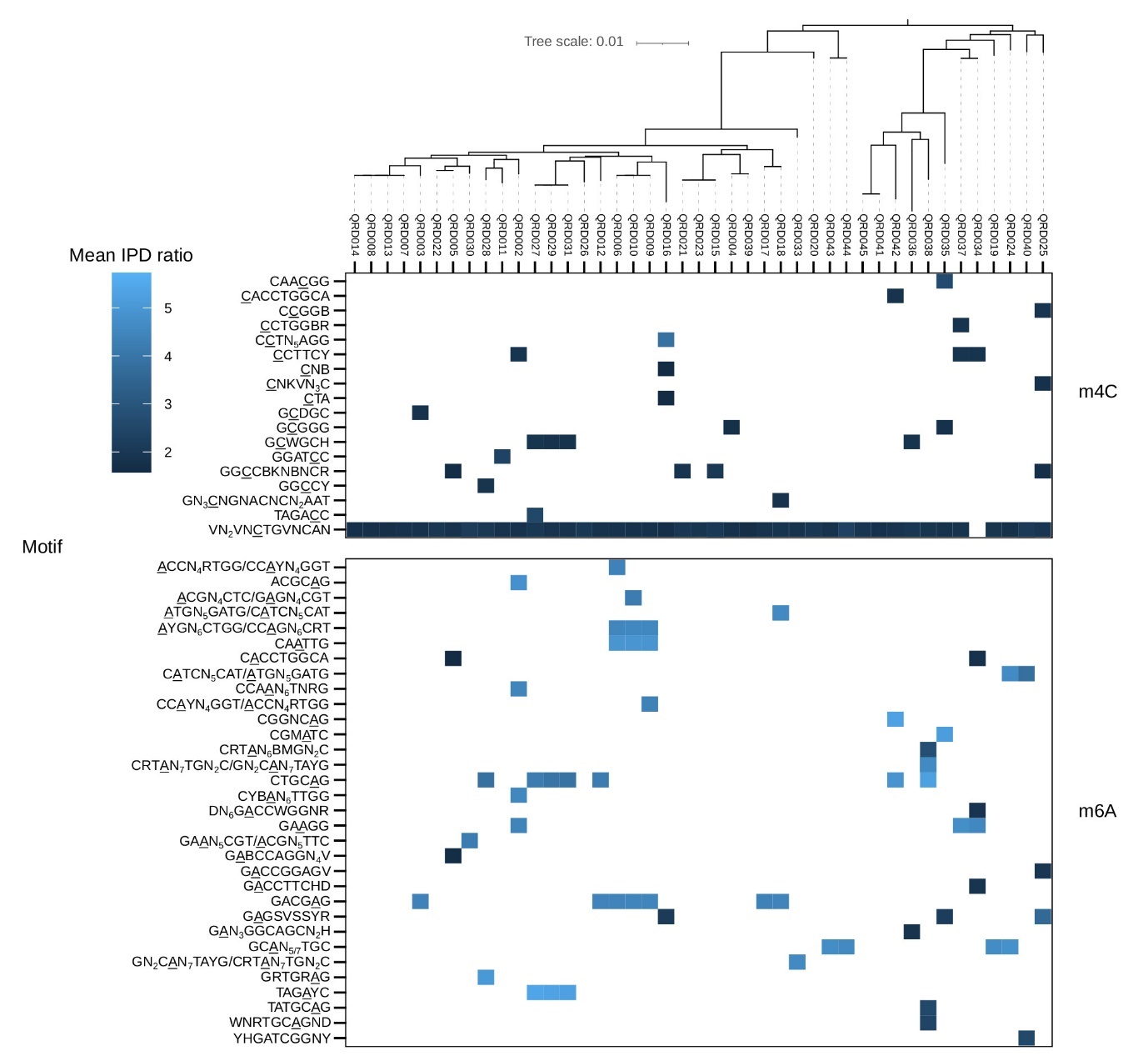

**Figure S4. DNA methylation motifs are not associated with phylogenetic clustering, related to Figure 2.**

DNA methylation motifs as annotated by the PacBio CCS sequencing platform and maximum likelihood core genome phylogenetic tree. Motifs that are present are indicated by their respective mean interpulse duration (IPD) ratio. Similar or identical motifs were grouped manually for simpler viewing. Methylated bases are underlined.

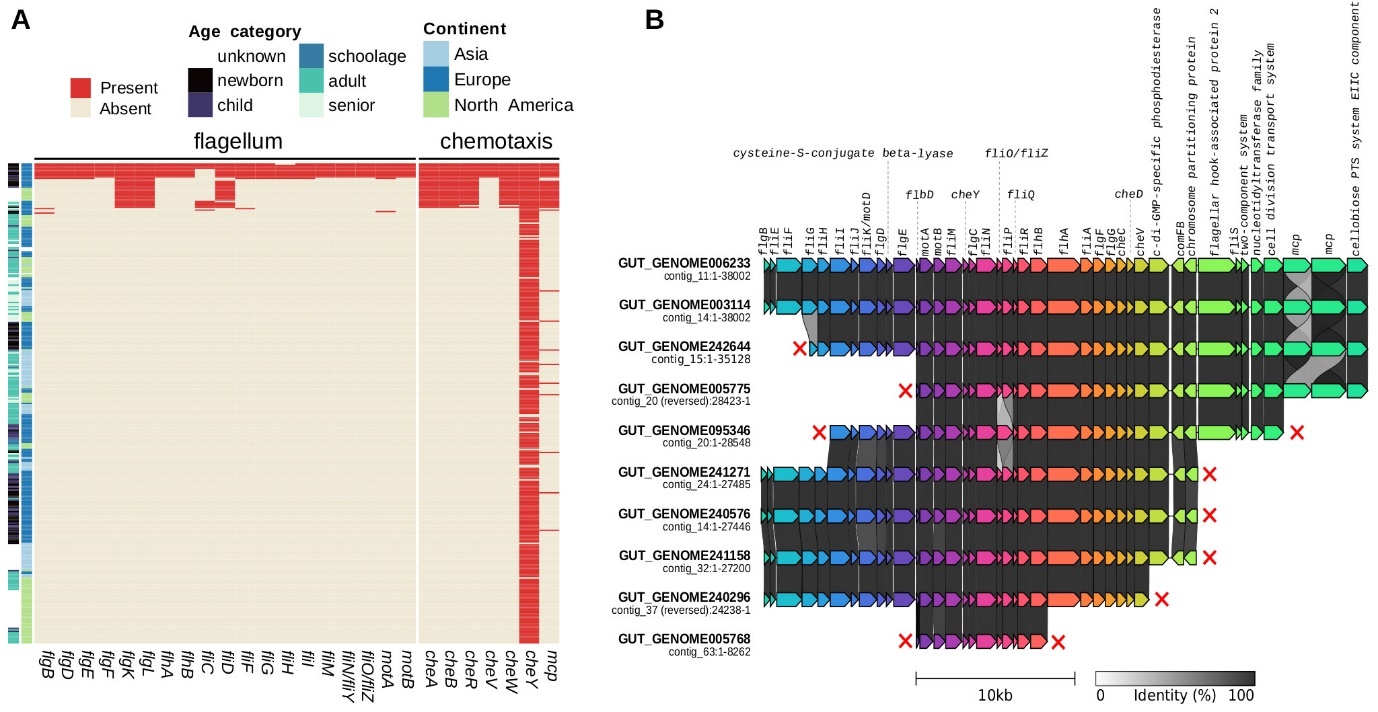

**Figure S5. Motility possibly restricted to strains from infants, related to Figure 2.**

(A) We annotated all genomes functionally using KEGG orthologs (KO) and pathways. A striking correlation was found with age and flagellum biosynthesis and chemotaxis. Presence and absence of essential flagellum and chemotaxis genes is indicated for each genome, annotated with age category and continent.

(B) Comparison of flagellum operon as detected in ten MAGs. Red crosses indicate contig ends (i.e., technical limitations rather than gene loss).

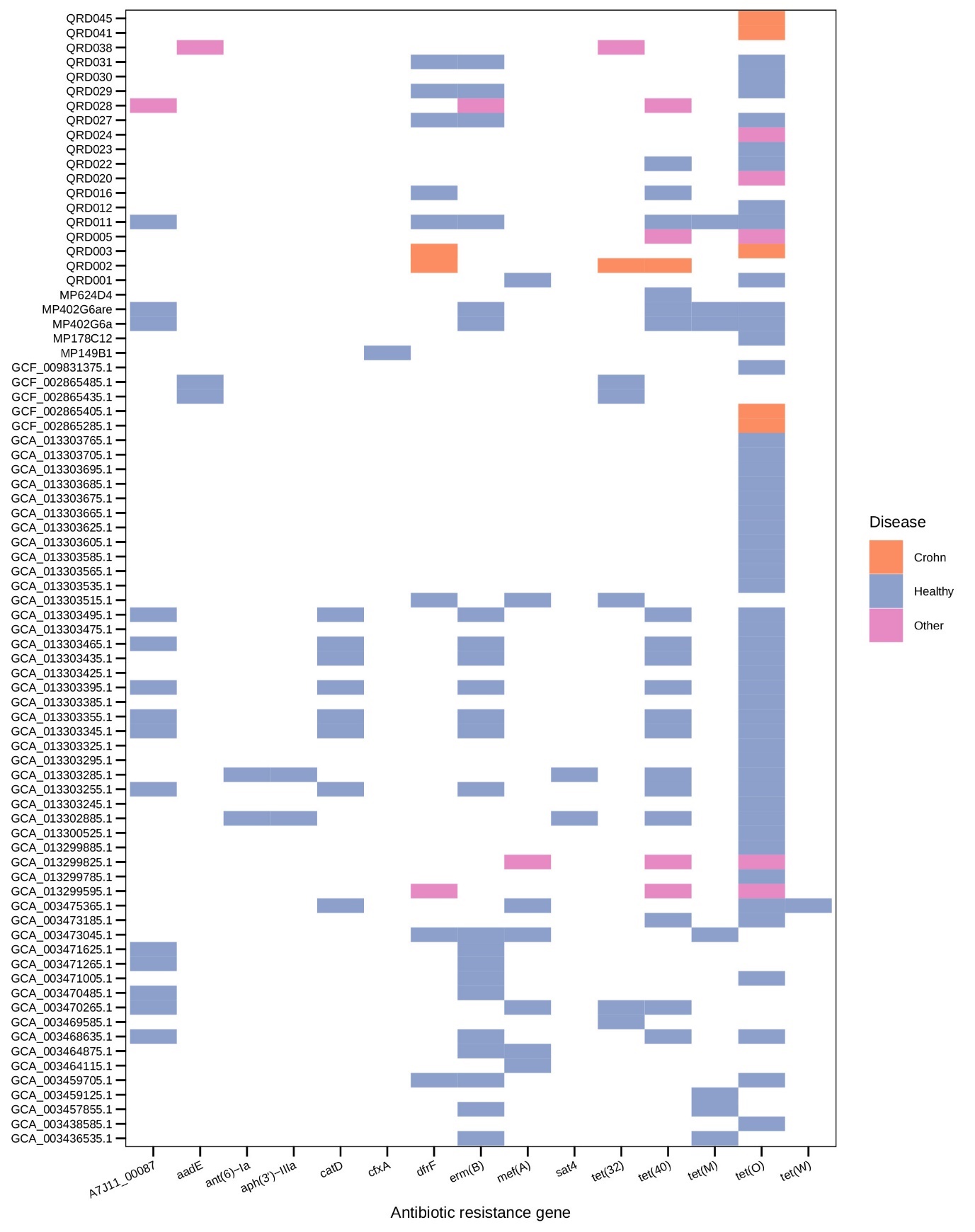

**Figure S6. Antibiotic resistance genes in *Ruminococcus* *gnavus* isolate genomes, related to figure 2.**

We screened all 125 isolate genomes for the presence of antibiotic resistance genes (92 derive from healthy people, 25 from Crohn’s disease patients and 8 from other diseases). Eighty-seven (62% of isolates) were found to have at least one resistance gene. Each gene hit is annotated with the host disease phenotype of the corresponding isolate.

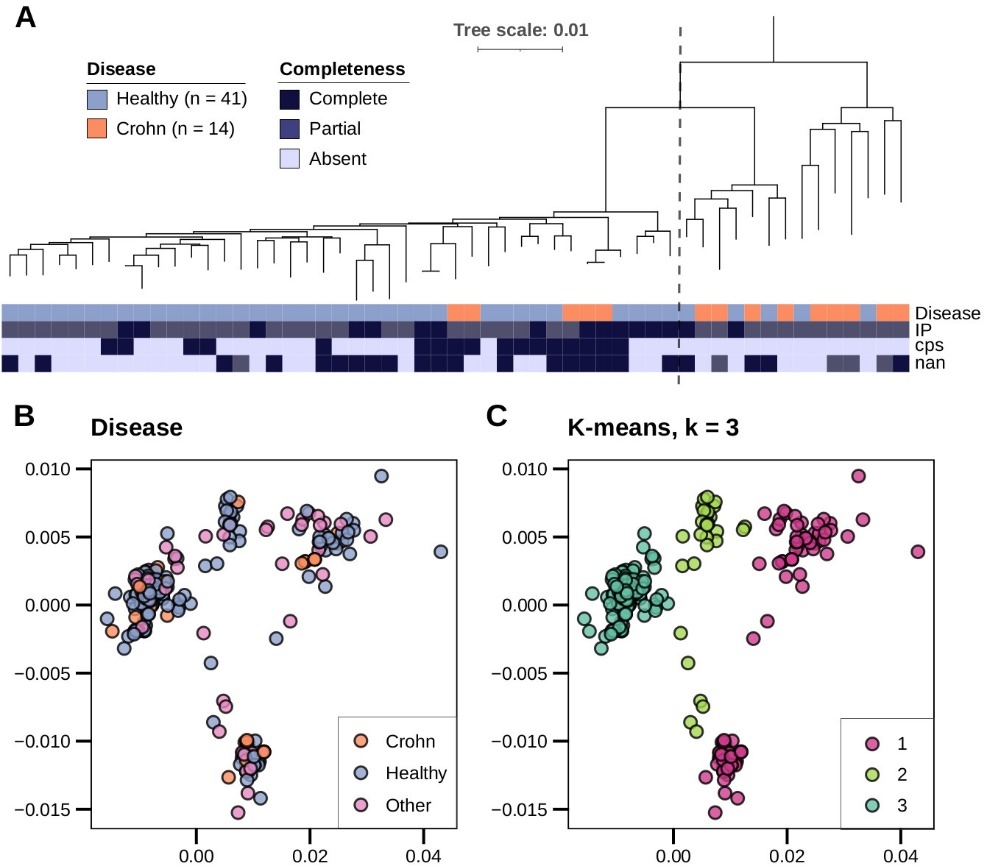

**Figure S7. Phylogenetic clades associated with host disease phenotype and capsule genes, related to Figure 3.**

(A) Maximum likelihood phylogenetic tree of core genomes of deduplicated isolates, annotated with host disease phenotype and putative virulence gene clusters that differ between isolates. The dashed line indicates the separation between the healthy-associated group and Crohn’s-associated group. The tree was midpoint rooted. IP: inflammatory polysaccharide (23 genes, ‘partial’ = 20 or 21 genes), cps: capsular polysaccharide (20 genes), nan: sialic acid metabolic cluster (11 genes, ‘partial’ = 6 genes).

(B) Principal Coordinate Analysis (PCoA) of core genome phylogenetic distances of all 333 *R. gnavus* genomes. The first figure shows each genome annotated by the disease state of the host, the second shows the same genomes clustered using k-means clustering (k = 3). Cluster 1 (magenta) contains 21 Crohn’s-derived genomes (orange) and 42 healthy-derived genomes (blue); cluster 3 (sea green) has 13 Crohn’s genomes and 153 healthy genomes.

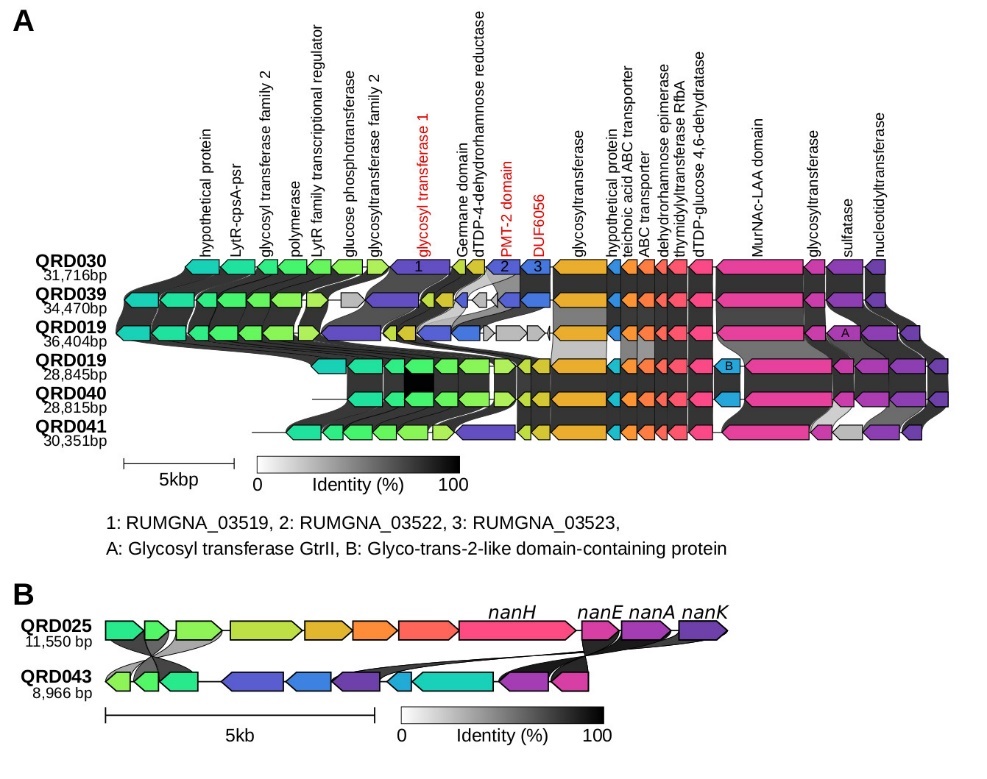

**Figure S8. Genomic architecture of operons differs between strains, related to Figure 3.**

(A) We identified six genes or gene clusters in literature that were putatively associated with gut inflammation or a Crohn’s disease phenotype, see also Figure 3A. Characterisation of these gene clusters in complete *R. gnavus* genomes showed that two clusters, encoding an inflammatory glucorhamnan polysaccharide and sialic acid metabolism, exist in two different varieties. Comparison of variations of the inflammatory polysaccharide operon as reconstructed from PacBio genomes. The top gene cluster is annotated with functions as predicted by Bakta. Genes absent from the alternative cluster are marked with red annotation and numbers 1-3. Genes inserted in alternative clusters are labelled A and B.

(B) Comparison of variants of the *nan* operon for sialic acid metabolism. Previously described *nan* genes are annotated.

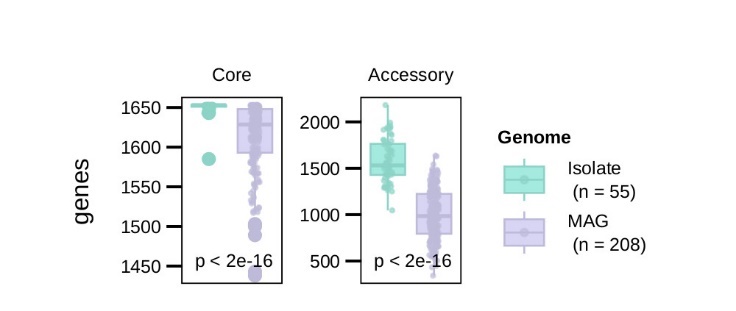

**Figure S9. Isolate genomes are more complete and suitable for GWAS than MAGs, related to Figure 3.**

Comparison of core and accessory genome size between genomes derived from isolates or metagenome-assembled genomes (MAG). P-values calculated using Wilcoxon rank sum test.

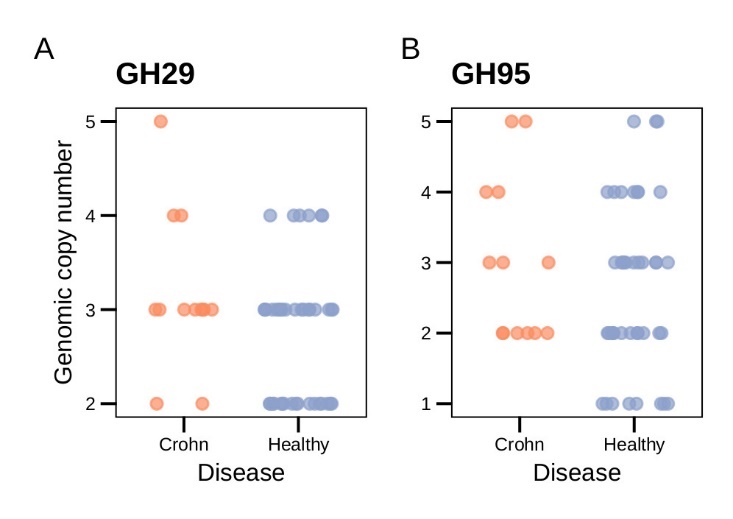

**Figure S10. Fucosidase CAZyme families not different in isolates from healthy people or Crohn’s patients, related to Figure 3.**

(A) Genomic copy numbers of CAZyme families encoding fucosidases involved in mucin degradation compared between isolates derived from Crohn’s disease patients and healthy people. Glycosyl hydrolase family 29. Wilcoxon rank sum test, p = 0.098.

(B) Glycosyl hydrolase family 95. Wilcoxon rank sum test, p = 0.39.
